# Endosome maturation is orchestrated by inside-out proton signaling through a Na^+^/H^+^ exchanger and pH-dependent Rab GTPase cycling

**DOI:** 10.1101/2024.12.09.627558

**Authors:** YouJin Lee, Qing Ouyang, Hasib Aamir Riaz, Li Ma, Morgan Fleishman, Michael Schmidt, Jeffrey L. Dupree, David G. Lambright, Eric M. Morrow

## Abstract

Endosome maturation requires progressive lumen acidification. To what extent is lumen acidification sensed by cytosolic-side molecules that drive endosome maturation? We show here that “inside-out” proton signaling through the endosomal Na^+^/H^+^ Exchanger 6 (NHE6) activates the late endosome master regulator Rab7. The mechanism involves potent inactivation of the Rab7 GTPase-activating protein (GAP) TBC1D5 with decreasing pH. NHE6 interacts with TBC1D5 in a complex with Rab7. Neurons from NHE6-null mice or mice engineered with a selective defect in NHE6 proton efflux exhibit blocked endosome maturation and decreased active Rab7, consistent with an overactive Rab7 GAP. Finally, epistatic knock-down of TBC1D5, thereby reducing Rab7 GAP activity, in NHE6-null neurons rescues Rab7 GTPase cycling and endosome maturation. Importantly, NHE6 is mutated in Christianson Syndrome underscoring the significance of these mechanisms to neurodegeneration. We conclude that lumen acidification regulates pH-dependent Rab GTPase cycling to coordinate late endosome maturation by a process involving proton signaling.

## Introduction

Endosomes mature from early to late endosomes, and eventually fuse with lysosomes^1^. Endosome maturation involves coordinated regulation of Rab protein activity on cytosolic-side, membrane surfaces, as well as progressive acidification of the endosomal lumen^1,2,3,4^. Rab proteins are small Ras-like GTPases, that are highly conserved from yeasts to mammals^5,6^, and cycle between active GTP-bound and inactive GDP-bound states^7,8,9,10,11^. During trafficking, each intra-cellular compartment is associated with distinct Rab proteins and their effectors that control downstream functions. For example, Rab7 is localized to late endosomes and regulates a variety of late endosome functions, including trafficking and endosome-lysosome fusion^4,9^. Active GTP-bound Rab7 binds to effectors, such as Rab Interacting Lysosome Protein (RILP) on endosomal membranes^12,13^. Active GTP-bound Rab7 is deactivated by the Rab7 GTPase-activating protein (GAP) TBC1D5^14,15^. Once deactivated, inactive GDP-bound Rab7 no longer binds to these effectors and dissociates from the endosome membrane^16,17,18,19^_._

In addition to cycling of Rab GTPases, endosome maturation is also associated with gradual acidification of the endosome lumen^2,20^. In neurons, the endosomal lumen becomes progressively more acidic as endosomes mature and move retrograde from the distal projections toward the soma where they may fuse with lysosomes^13,21,22,23,24,25,26,27,28^. This acidification is mediated by the vacuolar ATPase (v-ATPase), which actively pumps protons into endosomes^20,29,30^. By contrast, endosomal Na^+^/H^+^ exchangers (NHEs), such as NHE6 which is abundant on early and late endosomes, permit proton efflux from endosomes. Long-standing questions in the field of endosome biology are: To what extent is endosome maturation driven by lumen acidification? What are the mechanisms whereby the cytosolic machinery governing endosome maturation senses lumen acidification? In this study, we demonstrate that endosomal NHE6 functions in “inside-out” proton signaling, thereby linking lumen acidification with Rab GTPase activity.

Importantly, loss-of-function mutations in NHE6 cause the X-linked neurological disorder Christianson Syndrome (CS). CS is characterized by postnatal microcephaly, intellectual disability, non-verbal status with autistic features, and involves neurodegeneration with motor abnormalities^31,32,33^. In our previous study, we generated an NHE6-null rat which exhibits early evidence of endosome and lysosome dysfunction, preceding prominent axonal loss^34^. However, despite the broad relevance of endolysosomal processes in both neurodevelopment and neurodegeneration^35,36,37^, the molecular mechanisms mediating pathogenesis in CS remain unclear. Our laboratory and others have shown, in variety of cell types, that loss of NHE6 leads to over-acidified endosomes^27,38,39,40,41^, suggesting that NHE6 plays a role in regulation of intra-endosomal pH; however, the endogenous functions of endosomal NHEs remain poorly defined. We have previously demonstrated that NHE6-null neurons exhibit defects in endosome maturation, including impairments in endosome-lysosome fusion^21,27^. This endosome-lysosome fusion defect is conserved in the homologous yeast mutant Nhx1^42^.

In this study, we define a new function for endosomal NHEs in inside-out proton signaling, linking endosome lumen acidification to Rab GTP-GDP cycling. We discover that the Rab7 GAP, TBC1D5, acts as a pH-sensor that detects and is inactivated by proton efflux through NHE6. In the absence of NHE6, the Rab7 GAP is overactive, leading to decreases in active GTP-bound Rab7, and impaired late endosome trafficking and endosome-lysosome fusion. To define the specific role of NHE6- mediated proton efflux in endosome maturation, we generated an efflux-defective NHE6 (NHE6-ED) mutant mouse. In neurons from this mouse, we find that Rab7 GTP-GDP cycling requires proton efflux through NHE6 for endosomal maturation. We also demonstrate that NHE6 and the Rab7 GAP physically interact and co-localize with Rab7 on late endosomes. Furthermore, knock-down of TBC1D5 in NHE6-null neurons rescues endosome trafficking, Rab7 GTP-GDP cycling, and endosome-lysosome fusion defects. In summary, we reveal a new function of the CS protein NHE6 as a signaling molecule, using protons to drive endosome maturation. Furthermore, our studies identify inside-out proton signaling as a mechanism regulating pH-dependent cytosolic molecules, and thereby orchestrating critical endosome processes.

## Results

### Loss of NHE6 leads to enlarged Rab7-positive late endosomes and trafficking defects

NHE6 is abundantly expressed on endosomes in axons and dendrites^27,43^. In our previous study, we observed axonal pathology and loss in aged NHE6-null rat brains at 12 months^34^. To investigate the impact of loss of NHE6 on neuronal processes preceding degeneration, we stained coronal sections from the corpus callosum and cortex of NHE6-null rats and their littermate controls using an SMI32 antibody. SMI32 is known to indicate axonal damage by recognizing non-phosphorylated neurofilament heavy (NF-H) existing in neuronal axons^44,45,46^. At 2 months, we observed an increased average size of SMI32 punctate and a greater number of enlarged SMI32-stained puncta in the corpus callosum (CC) and cortex (CTX) in NHE6-null rats compared to those of wild-type (WT) rats (Fig. 1a). Over time, the intensity of SMI32 staining increased in NHE6-null rats compared to WT rats (Extended Data Fig. 1a). In addition, axonal degeneration was confirmed in NHE6-null rat brains at 2 months using electron microcopy (Extended Data Fig. 1b).

**Fig. 1:**
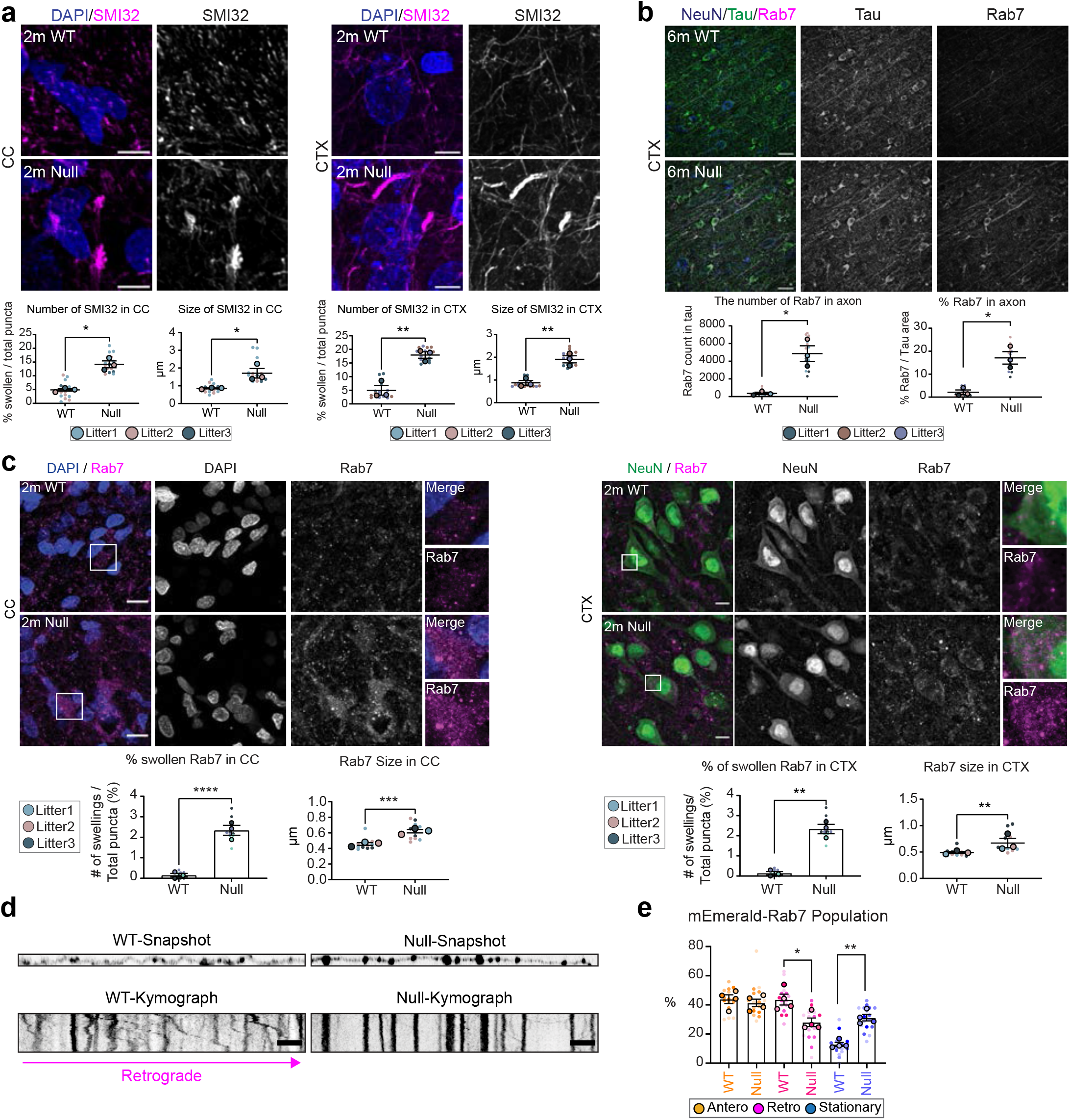
Rab7 late endosome defects in NHE6-null rats. **a,** Enlarged SMI32-stained (axonal damage marker, magenta) swellings in the corpus callosum (CC) and cortex (CTX) of NHE6-null rat brains at 2 months. The number of SMI32-stained swellings were divided by the total number of SMI32 puncta along with the average size of puncta. (a-c) Means from each independent experiment (big dots, WT = 3, Null = 3 animals) overlay the entire dataset (small dots, 5 sections per each animal) and used for statistical analysis. Animals from same litters are color-coded. Two-tailed unpaired t-test with Welch’s correction. Scale bar, 5 μm. **b,** Accumulation of enlarged Rab7 late endosomes in the axons of NHE6-null rat cortex at 6 months. The number of Rab7 puncta and % of Rab7 area covered were quantified within Tau-stained axons. Two-tailed unpaired t-test with Welch’s correction. Scale bar, 20 μm. **c,** Enlarged Rab7 late endosomes in NHE6-null rats at 2 months. The number of enlarged Rab7 was divided by the total number of Rab7 puncta (%) along with the average size of puncta. Two-tailed unpaired t-test with Welch’s correction. Scale bar, 10 μm. **d,** Representative snapshots and kymographs from WT and NHE6-null neurons show the movement of mEmerald-Rab7. Retrograde direction is indicated. Scale bar, 1 μm. **e,** Increased numbers of stationary mEmerlad-Rab7 labelled endosomes in NHE6-null neurons. Decreased numbers of retrograde Rab7 endosomes were also detected in NHE6-null neurons. Anterograde, retrograde, and stationary endosomes were divided by the total number of mEmerlad-Rab7 endosomes. Means from each animal (big dots, WT = 4, Null = 4 pups) overlay the entire dataset (small dots, n = 17-20 neurons from four independent experiments) and used for statistical analysis. Ordinary one-way ANOVA with Tukey’s HSD. Data are represented as mean ± SEM. ****p < 0.0001, ***p < 0.001, **p < 0.01, and *p < 0.05.

We previously observed the accumulation of late endosomes in NHE6-null neurons and brains prior to axonal degeneration^21,34^. To further evaluate the extent to which late endosomes accumulated in swollen axons, we stained the cortex sections from WT and NHE6-null rats using Rab7 (a late endosomal marker) and Tau (an axonal marker) antibodies. We observed an increased number and size of Rab7-positive puncta in the axons in NHE6-null rat brains compared to WT (Fig. 1b). The number of swollen Rab7- positive endosomes (diameter larger than 1.5 μm) and average size also were quantified in the corpus callosum and cortex sections from WT and NHE6-null rat brains at 2 months (Fig. 1c). NHE6-null rats exhibited both a higher number and larger size of Rab7-positive structures in the corpus callosum and cortex when compared to WT rats. As NHE6 is also localized on early endosomes^27,39,47^, we also stained sections from WT and NHE6-null rat brains with Rab5 (early endosomal marker). However, we did not detect swollen Rab5-positive endosomes or significant differences in the average size of Rab5-positive endosomes between WT and NHE6-null rats (Extended Data Fig. 2a).

Previous studies have linked axonal loss to endolysosomal trafficking defects^44,48,49^. To investigate Rab7 trafficking, we transfected mEmerald-Rab7 into primary neurons from WT and NHE6-null rats (Fig. 1d). We confirmed that Rab7 endosomes bidirectionally moved in anterograde and directions (Fig. 1d)^50^. However, NHE6-null neurons have a higher number of stationary mEmerald-Rab7 and fewer retrogradely moving mEmerald-Rab7 endosomes in comparison to WT neurons (Fig. 2c). Also, the speed of both anterograde and retrograde Rab7 was faster in NHE6-null neurons than in WT neurons (Extended Data Fig. 3a). Furthermore, the size of mEmerald-Rab7 (Extended Data Fig. 3b, c, d) and endogenous Rab7-positive endosomes (Extended Data Fig. 3e) is larger in primary NHE6-null neurons compared to those in WT neurons. By contrast, we did not observe trafficking defects in mEmerald-Rab5 labelled endosomes in NHE6-null neurons (Extended Data Fig. 2b, c). In summary, our findings indicated that NHE6-null neurons exhibited an accumulation of enlarged Rab7 endosomes and late endosomal trafficking defects.

**Fig. 2:**
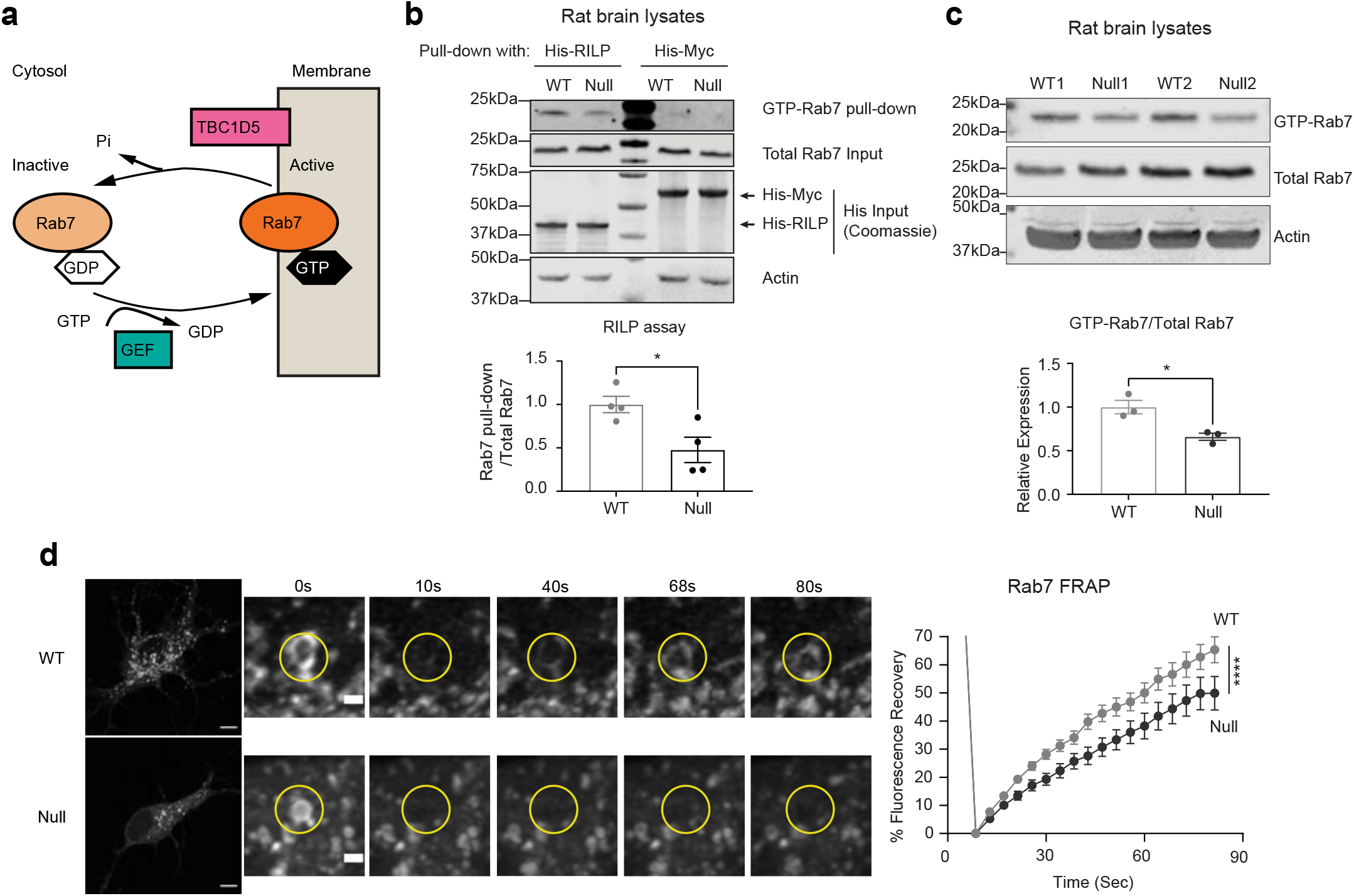
Active GTP-bound Rab7 is decreased in NHE6-null rat brains. **a,** Schematic of GTP-GDP exchange cycle of Rab7. TBC1D5 is an GTPase activating protein (GAP) of Rab7. **b,** Reduced RILP-bound Rab7 (GTP-Rab7 pull-down) in NHE6-null rat brains at 2 months. Lysates from NHE6-null and WT littermate rat brains were incubated with His- RILP recombinant proteins or His-Myc (negative control) for the pull-down assay. Once samples were eluted, Rab7 was detected by western blot. The densitometry of GTP- Rab7 was divided by the total Rab7 for the quantification (4 rat brains for each genotype). Two-tailed unpaired t-test with Welch’s correction. **c,** Decreased GTP-bound Rab7 in NHE6-null rat brains at 2 months. Lysates from WT and NHE6-null littermate rat brains were incubated with GTP agarose to enrich GTP- bound protein fractions. Rab7 was detected in the GTP-enriched fraction by western blot. The densitometry of GTP-bound Rab7 was divided by the total Rab7 for the quantification (3 rat brains for each genotype). Two-tailed unpaired t-test with Welch’s correction was performed. **d,** Representative images of fluorescence recovery after photobleaching (FRAP) assay with mEmerlad-Rab7-transfected primary neurons at different time points. Slower recovery of mEmerlad-Rab7 was detected from primary NHE6-null neurons compared to WT littermate neurons. Yellow circles indicate where photobleaching occurs and the fluorescence intensity was measured (WT = 4, Null = 4 pups, n = 35-37 neurons from four independent experiments). Repeated measure two-way ANOVA with Sidak’s post hoc. Scale bar, 5 μm. Scale bar for time course images, 0.1 μm. Data are represented as mean ± SEM. ****p < 0.0001, ***p < 0.001, **p < 0.01, and *p < 0.05.

### Active GTP-bound Rab7 is decreased in NHE6-null neurons

Rab7 is a small GTPase that cycles between active GTP-bound and inactive GDP-bound states (Fig. 2a)^7,8,9,10,11^. The GTP-GDP cycle of Rab7 is required for late endosomal maturation and retrograde trafficking in neurons^13,51^. As more stationary Rab7 endosomes were detected in NHE6-null neurons (Fig. 1d, e), we hypothesized that defective GTP-GDP cycling of Rab7 disrupts the retrograde transport and leads to more stationary Rab7 endosomes. We measured the amount of active Rab7 using multiple approaches. We first performed a RILP pull-down assay (Fig. 2b), as active GTP-bound Rab7 binds to RILP^52,53^. We incubated lysates from WT and NHE6-null rat brains at 2 months with His-RILP probes. A lower amount of active Rab7, bound to His- RILP probes in the lysates, was detected in NHE6-null rat brains compared to those of WT brains. We did not observe changes in the total Rab7 expression between WT and NHE6-null rat brains. To confirm our findings on reduced active GTP-Rab7, we also performed a GTP-agarose pull-down assay (Fig. 2c)^54,55^. The lysates from WT and NHE6-null rat brains were incubated with a GTP-agarose to enrich the pool of GTP-bound proteins and followed by detection with anti-Rab7 antibody. A lower amount of active Rab7 bound to GTP-agarose was found in NHE6-null rat lysates compared to WT rat lysates. Notably, regarding these two assays, we did not observe a significant difference in the amount of active Rab5 bound to His-RILP (Extended Data Fig. 2d), or active Rab5 bound to GTP-agarose (Extended Data Fig. 2e) in WT as compared to NHE6-null rats. Lastly, we performed fluorescence recovery after photobleaching (FRAP) in primary WT and NHE6-null neurons transfected with mEmerland-Rab7 (Fig. 2d). We bleached a small region of Rab7 in the soma and recorded the fluorescence recovery over time. Recovery of fluorescence occurs when photobleached mEmerald- Rab7 diffuses from membrane and is replaced by fresh mEmerald-Rab7 protein or is delivered via RabGDI from the cytosol upon GTP-GDP exchange cycle^56^. Previous studies showed that the GTP-GDP status of Rab7 affected its fluorescence recovery^57,58^. The recovery of mEmerald-Rab7 in primary WT neurons reached to ∼60 % within 80 seconds (Fig. 2d). In contrast, the recovery of mEmerlad-Rab7 in primary NHE6-null neurons was slower and reached to ∼45 % over the same time course. Overall, these corroborative results (from the RILP assay, GTP-agarose pull-down assay, and the FRAP studies) indicate decreased active GTP-bound Rab7 in NHE6-null neurons.

### Increased co-localization of Rab7 with its GTPase-activating protein TBC1D5 in NHE6-null neurons

Given our finding that the GTP-GDP cycling of Rab7 is disrupted in NHE6-null neurons (Fig. 2), we hypothesized that this is due to disrupted regulation of Rab7 activity by its GTPase-activating protein (GAP), TBC1D5^14,15^. Previous studies have suggested that Nhx1, the NHE6 homologue in yeast, interacts with Gyp6, the TBC1D5 orthologue^42,59^. TBC1D5 is localized to late endosomes^15,53,60^, regulating GTP hydrolysis of GTP-bound Rab7 (as schematized in Fig. 2a). First, we tested whether TBC1D5 acts as a Rab7-specific GAP^14,15,61^. We measured Rab7 GTPase and Rab5 GTPase activities following incubation with varying concentrations of TBC1D5. TBC1D5 accelerates the GTP hydrolysis of Rab7, and consequently, more phosphates will be released during this process. As expected, with increasing concentrations of TBC1D5, more GTP hydrolysis of Rab7 was detected (Fig. 3a). However, TBC1D5 did not effectively hydrolyze Rab5 (Extended Data Fig. 2f).

**Fig. 3:**
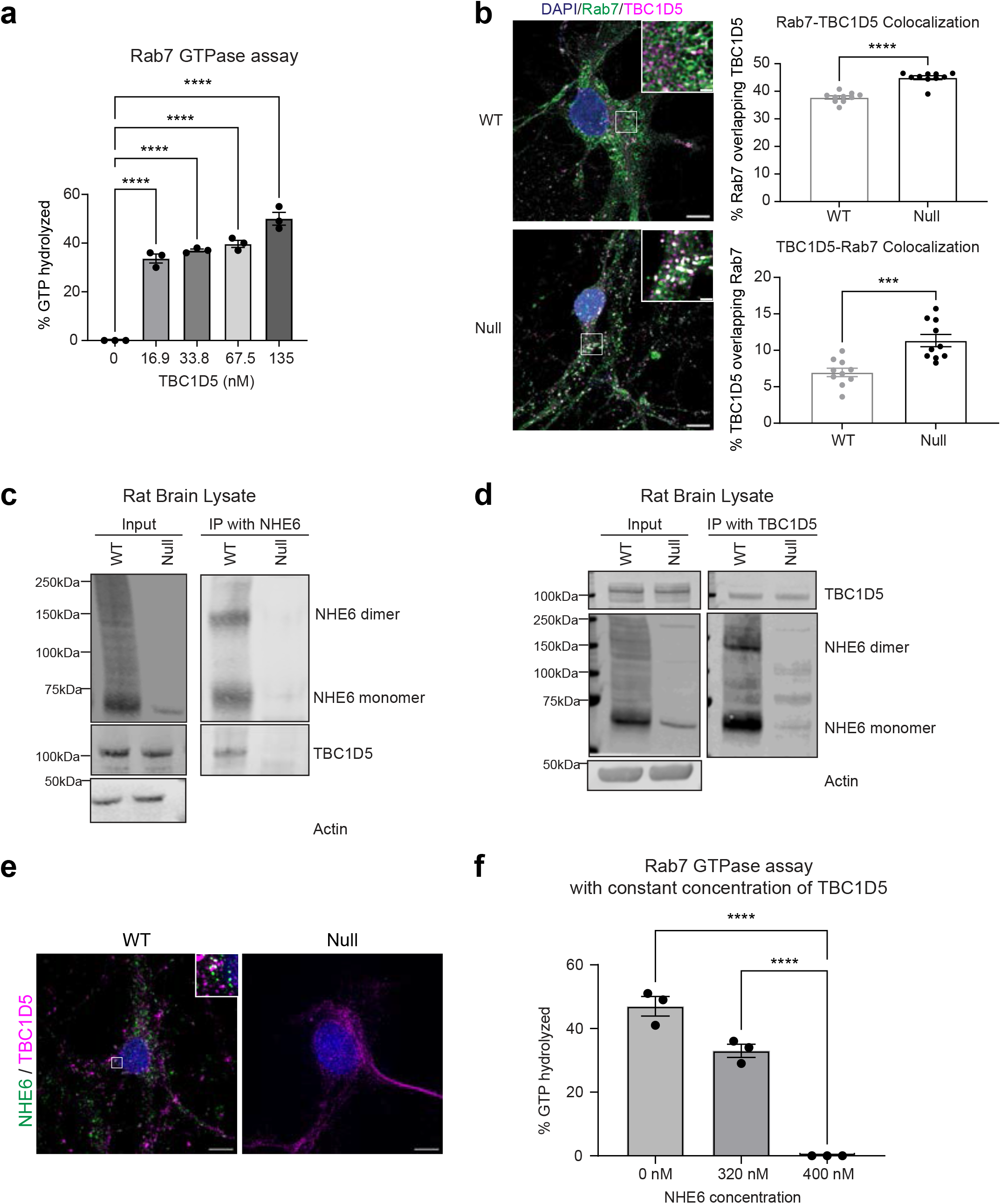
The interaction of NHE6 and TBC1D5 affects the Rab7 GTPase activity. **a,** *In vitro* Rab7 GTPase assay shows that higher concentrations of TBC1D5 increased GTP hydrolysis of Rab7 measured by phosphate releases. Pre-loaded GTP-Rab7 was incubated with various concentration of recombinant TBC1D5 for the luminescence measurement. Ordinary one-way ANOVA with Tukey’s HSD (n = 3 independent experiments). **b,** Increased co-localization of Rab7 (green) and TBC1D5 (magenta) in NHE6-null neurons. The quantifications show the Mander’s coefficient (M1: Rab7 overlapping TBC1D5, M2: TBC1D5 overlapping Rab7) from high-content imaging. Each dot indicates a mean of each pup (WT = 10, Null = 10 pups, n = 10,000-14,000 neurons imaged from each pup). Two-tailed unpaired t-test with Welch’s correction. **c,** Co-immunoprecipitation of NHE6 with TBC1D5 from WT and NHE6-null (negative control) rat brains at 2 months. **d,** Reciprocal immunoprecipitation from WT and NHE6-null littermate rat brain lysates at 2 months shows co-immunoprecipitation of TBC1D5 with NHE6. **e,** Representative images of primary rat neurons show the co-localization of NHE6 (green) and TBC1D5 (magenta). Primary neurons were imaged using structured illumination microscopy. NHE6-null neurons were used as a negative control. Scale bar, 5 μm. **f,** *In vitro* Rab7 GTPase assay with increasing concentrations of NHE6. GTP-Rab7 and constant concentration of TBC1D5 proteins were incubated with increasing concentration of NHE6 proteins to measure phosphate release. Higher concentrations of NHE6 decreased GTP hydrolysis of Rab7 mediated by TBC1D5. Ordinary one-way ANOVA with Tukey’s HSD. Three independent experiments were conducted. Data are represented as mean ± SEM. ****p < 0.0001, ***p < 0.001, **p < 0.01, and *p < 0.05.

We next investigated the localization of Rab7 and TBC1D5 in primary neurons from WT and NHE6-null rats using high-resolution microscopy with the Nikon Spatial Array Confocal (NSPARC) detector (Fig. 3b). Notably, NHE6-null neurons exhibited a markedly increased co-localization between Rab7 and TBC1D5 compared to WT neurons. This supports the model that NHE6 in the late endosome membrane limits TBC1D5 action on Rab7, and in the absence of NHE6, there is dysregulation of TBC1D5 activity on Rab7, leading to less active GTP-bound Rab7.

### NHE6 interacts with the Rab7 GAP TBC1D5 at the endosome membrane

To investigate whether NHE6 interacts with TBC1D5 in neurons, reciprocal co-immunoprecipitation assays were conducted in lysates from both WT and NHE6-null rat brains at 2 months. Specifically, the lysates were immunoprecipitated with an anti-NHE6 antibody, and TBC1D5 was detected in the precipitants from WT rat brains (Fig. 3c).

Reciprocal immunoprecipitations were also carried out using an anti-TBC1D5 antibody, confirming the TBC1D5 interaction with NHE6 (Fig. 3d). We also observed the co- localization of NHE6 and TBC1D5 in primary WT neurons using structure illumination microscopy (SIM) super-resolution microscopy (Fig. 3e).

Next, we examined the extent to which NHE6 protein interaction with TBC1D5 affects TBC1D5-mediated Rab7 GTPase activity *in vitro* (Fig. 3f). Recombinant Rab7 and TBC1D5 proteins were incubated with increasing concentrations of NHE6. Our data revealed that increasing concentration of NHE6 decreased the GTP hydrolysis of Rab7. These findings suggest that NHE6 biochemically hindered TBC1D5-mediated GTP hydrolysis of Rab7. Overall, our data suggest that the interaction between NHE6 and TBC1D5 limits TBC1D5 activity on Rab7.

### The proton efflux function of NHE6 regulates Rab7 activity

We next asked if proton efflux via NHE6 regulates Rab7 activity. To test this, we used CRISPR/Cas9 genome editing to generate highly specific point mutations in NHE6 (NHE6-ED) mice, wherein the exchanger function of NHE6 is inactive. Two mutations (E255Q/D260N) in the cation exchanger domain of human NHE6, are known to impede ion-proton transport^27,39^. The corresponding glutamic acid and aspartic acid in mouse NHE6 are at positions 235 and 240, respectively (Extended Data Fig. 4a). We designed guide RNAs to substitute glutamic acid with glutamine (c.705G>C, p.E235Q) and aspartic acid with asparagine (c.720G>A and 722C>T, p.D240N) to mimic constructs previously studied extensively *in vitro* (Extended Data Fig. 4b)^27,62^. Sanger sequencing of genomic DNA isolated from tail biopsy showed successful genome editing in NHE6- ED mutant mice (Extended Data Fig. 4c), indicating that the desired substitution mutations had been generated in mouse *Slc9a6*.

We studied the NHE6-ED mouse thoroughly to establish that NHE6 protein is made and trafficked appropriately, and impaired in proton efflux. Western blotting from mouse brain lysates confirmed the expression of NHE6 protein, which is stably expressed to an equivalent level of both NHE6 monomer (∼70 kDa) and dimer forms (∼140 kDa) between WT and NHE6-ED mice (Extended Data Fig. 4d). Also, importantly, we measured the intra-endosomal pH in primary neurons from NHE6-ED and its littermate WT mice by fluorescent ratio imaging fluorescein isothiocyanate (FITC)-conjugated transferrin (Tfn; pH sensitive) to AlexaFluor-546-conjugated Tfn (pH insensitive) (Fig. 4a)^63^. Our data demonstrate decreased lumen pH in neuronal endosomes both in the some and neurites in NHE6-ED neurons as compared to WT neurons (Fig. 4b, c). This indicates compromised proton efflux function of the NHE6-ED mutant endosomes, akin to NHE6-null endosomes^27^. Primary neurons from WT and NHE6-ED mice were stained with Rab7 and NHE6, and no differences in the NHE6 colocalization with Rab7 were detected (Extended Data Fig. 4e). We conducted immunoprecipitation of NHE6 to confirm the interaction of NHE6-ED and TBC1D5 in the lysates from NHE6-ED mouse brains (Extended Data Fig. 4f), suggesting that the relevant protein structure of NHE6- ED was intact. Overall, given the extent of our control studies, the NHE6-ED mouse mutant appears to demonstrate proper NHE6 protein localization in endosomes with a precise defect in proton efflux, reflected by over-acidification of endosomes.

**Fig. 4:**
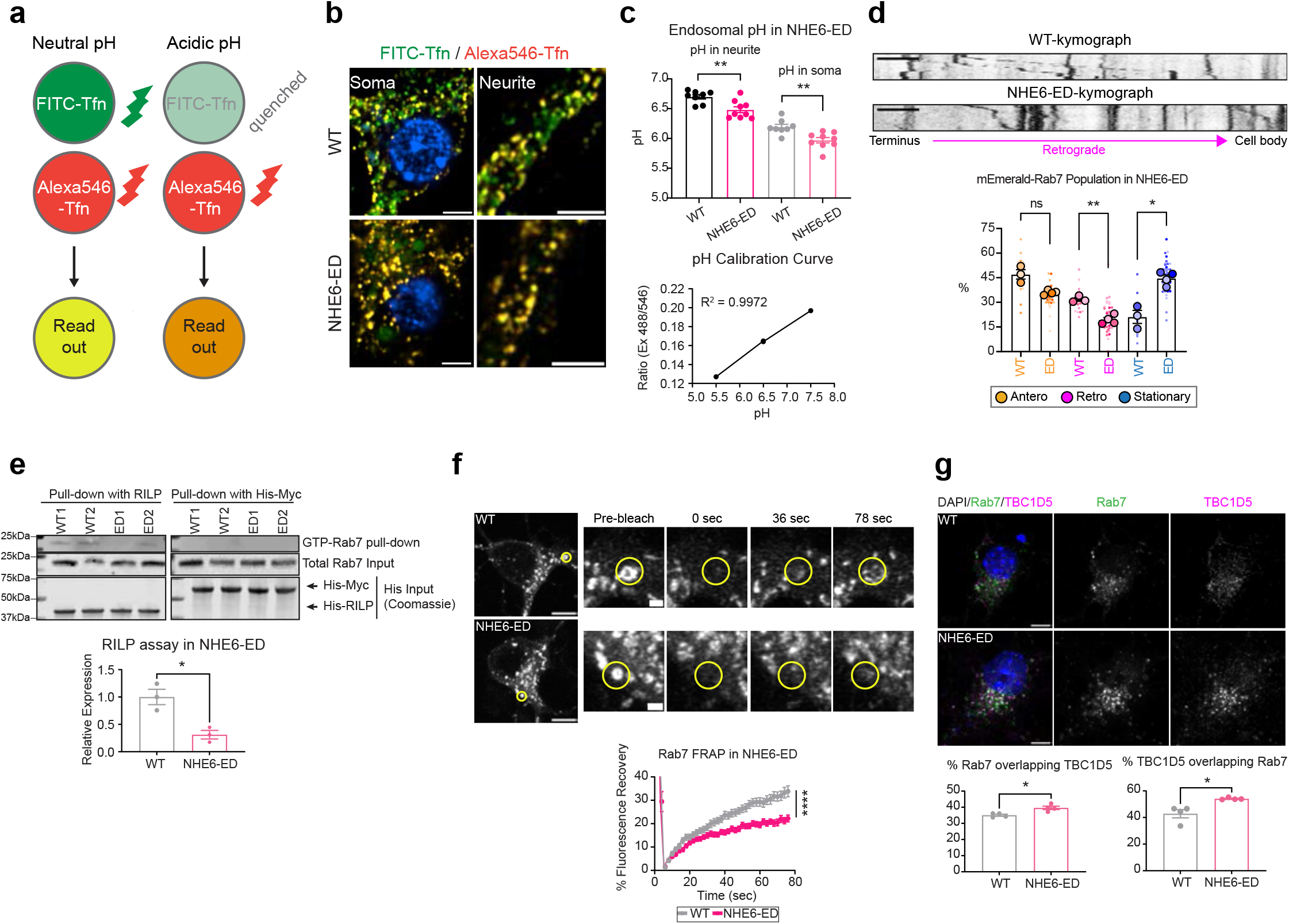
Regulation of Rab7 activity requires NHE6-mediated proton efflux. **a,** Schematic of the luminal pH measurement. Primary neurons are loaded with pH- sensitive, FITC-conjugated transferrin (Tfn) and pH-insensitive Alexa Fluor546-Tfn. More orange color indicates Tfn-positive endosomes with lower luminal pH. **b,** The decreased pH of endosomes in the neurites and soma of NHE6-ED neurons. Scale bar: 5um. **c,** Lower luminal pH of transferrin-positive endosomes in the neurites and soma of NHE6-ED neurons. The calibration curve was used to determine the luminal pH of transferrin-positive endosomes. Each dot indicates each pup (WT = 8, NHE6-ED = 9 pups, n = 10,000-14,000 neurons imaged from each pup). Two-tailed unpaired t-test with Welch’s correction. **d,** Representative kymographs show the movement of mEmerald-Rab7 endosomes. Retrograde direction is indicated. Scale bar, 2 μm. Increased numbers of stationary mEmerlad-Rab7 endosomes along with decreased retrograde numbers were detected in primary NHE6-ED neurons. Means from each animal (big dots, WT = 3, NHE6-ED = 4 pups) are overlay the entire dataset (small dots, n = 20-35 neurons from three independent experiments) and used for statistical analysis. Animals from same litters are color-coded. Ordinary one-way ANOVA with Tukey’s HSD. **e,** Decreased RILP-bound Rab7 (GTP-Rab7 pull-down) in NHE6-ED mouse brains at 2 months. Lysates from WT and NHE6-ED mouse brains were incubated with His-RILP recombinant proteins or His-Myc (negative control) for the pull-down assay. After elution, Rab7 was detected by western blot. The densitometry of GTP-Rab7 was divided by the total Rab7 (3 brains for each genotype). Two-tailed unpaired t-test with Welch’s correction. **f,** Slower FRAP recovery of mEmerlad-Rab7 in NHE6-ED neurons. Yellow circles indicate where photobleaching occurs and the fluorescence intensity was measured (WT = 3, Null = 4 pups, n = 30-47 neurons from three independent experiments). Repeated measure two-way ANOVA with Sidak’s post hoc. Scale bar, 5 μm. Scale bar for time course images, 0.1 μm. **g,** Increased co-localization of Rab7 (green) and TBC1D5 (magenta) in NHE6-ED neurons. Quantification shows the Mander’s coefficient (M1: Rab7 overlapping TBC1D5, M2: TBC1D5 overlapping Rab7) from high-content imaging. Each dot indicates a mean of each animal (WT = 4, Null = 4 pups, n = 10,000-14,000 neurons imaged from each pup). Two-tailed unpaired t-test with Welch’s correction. Scale bar, 5 μm. Data are represented as mean ± SEM. ****p < 0.0001, ***p < 0.001, **p < 0.01, and *p < 0.05.

We investigated endosome trafficking and maturation in NHE6-ED mouse neurons. We first investigated the extent to which trafficking of Rab7 endosomes is affected in NHE6-ED neurons. We monitored the movement of mEmerald-Rab7 endosomes in primary neurons from WT and NHE6-ED mice (Fig. 4d). NHE6-ED neurons have a higher number of stationary mEmerald-Rab7 endosomes and fewer retrograde mEmerald-Rab7 endosomes in comparison to WT neurons, akin to our prior observations in NHE6-null neurons (Fig. 1d). We did not observe differences in the speed of both anterograde and retrograde Rab7 endosomes (Extended Data Fig. 4g).

To determine the extent to which the proton efflux function of NHE6 affects GTP-GDP cycling of Rab7, we performed the RILP assay in WT and NHE6-ED brain lysates (Fig. 4e). Brain lysates were incubated with His-RILP and immunoblotted with anti-Rab7 antibody. A lower amount of active Rab7 bound to His-RILP probes in the lysates was detected in NHE6-ED mutant brains compared to those of WT brains, again akin to NHE6-null neurons. To further investigate the GTP-GDP exchange defects, we also performed the FRAP assay in primary WT and NHE6-ED neurons transfected with mEmerland-Rab7 (Fig. 4f). A small region of mEmerlad-Rab7 was photobleached and its recovery was recorded over time. The recovery of mEmerlad-Rab7 was slower in primary NHE6-ED neurons compared to those of primary WT neurons. Overall, neurons with the precise NHE6-ED mutation demonstrate similar defects to NHE6 null neurons.

Lastly, as increased co-localization of Rab7 with TBC1D5 was observed in NHE6- null neurons (Fig. 3b), we investigated the extent to which disrupted proton efflux caused by NHE6-ED affects TBC1D5 co-localization with Rab7. Primary WT and NHE6- ED neurons were stained with Rab7 and TBC1D5. The co-localization of Rab7 and TBC1D5 increased in NHE6-ED neurons compared to WT neurons (Fig. 4g), indicating that proton efflux via NHE6 is required for normal Rab7 interaction with TBC1D5 and that NHE6 proton efflux may reduce TBC1D5 activity on Rab7. In summary, our data show that proton efflux via NHE6 is required for proper Rab7 GTP-GDP cycling, regulating Rab7 localization with TBC1D5 and Rab7 activity.

### The GAP activity of TBC1D5 on Rab7 is pH dependent

Endosome maturation depends on acidification of the endosome lumen by the v-ATPase. We hypothesized that TBC1D5 acts as a sensor that detects acidification via proton efflux through NHE6 by adjusting its catalytic efficiency for acceleration of Rab7 GTP hydrolysis, i.e., converting GTP-bound to GDP-bound Rab7. In this way, NHE6 may function in inside-out signaling of endosome acidification, thereby promoting maturation by generating local decreases in pH on the cytoplasmic surface of the endosome.

In order to determine the catalytic efficiency of TBC1D5-catalyzed GTP hydrolysis on Rab7 over a range of pH values, we used well characterized differences in intrinsic tryptophan fluorescence between the GTP- and GDP-bound states of Rab7 to monitor the time course for GTP hydrolysis ^16^ in the absence and presence of the purified worm TBC1D5 orthologue (RBG-3, referred to as ceTBC1D5) (Fig. 5a, Extended Data Fig. 5a, and 5b). The expected increase in fluorescence intensity of Rab7 was observed as TBC1D5 accelerated GTP hydrolysis, which is followed by the rapid conformational switch from the GTP- to GDP-bound state. We determined the observed rate constant (kobs) at various concentrations of TBC1D5 (Fig. 5b and Extended Data Fig. 5c) under single turnover conditions using the purified TBC and GTPase domains of TBC1D5 and Rab7 as described previously^64^. The catalytic efficiency (kcat/KM) was determined from the slopes of linear fits to kobs as a function of TBC1D5 concentration. The observed kcat/KM depends strongly on pH and is well described by the simplest possible model, with active and inactive states, corresponding to titration of one or more groups with a common pKa (Fig. 5c). The intrinsic rate constant for Rab7 GTP hydrolysis, on the other hand, does not vary substantially over the same pH range in the absence of TBC1D5 (Extended Data Fig. 5a). These data indicate that the catalytic activity of TBC1D5 is strongly pH dependent, approaching full activity at the higher end of the pH range and minimal or no activity at the lower end. Moreover, the fitted pKa of 5.95 ± 0.02 is similar to the pKa of the sidechain of free histidine (∼6.04), suggesting histidine protonation state change may explain the pH sensitivity.

**Fig. 5:**
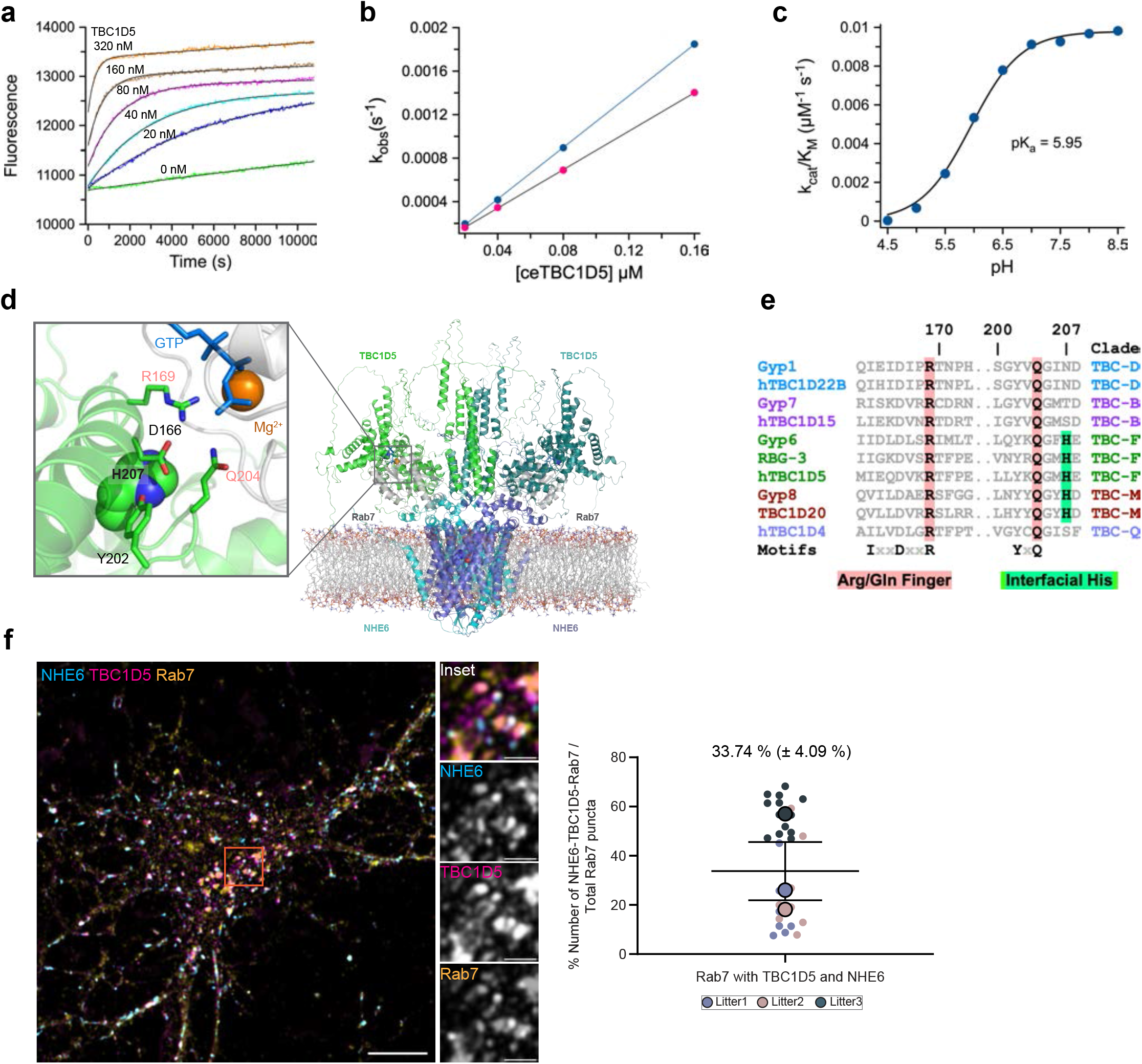
TBC1D5 activity is pH dependent. **a,** Intrinsic tryptophan fluorescence time courses of Rab7-GTP hydrolysis in the absence and presence of the *Caenorhabditis elegans* TBC1D5 ortholog (RBG-3, or ceTBC1D5) at pH 7.0. Different concentrations, as indicated, of ceTBC1D5 were added to purified Rab7-GTP *in vitro*. Solid lines are fits of exponential models to the experiment data. This assay was repeated in the pH range 4.5-8.5 to determine kobs at each pH value (Extended Data Fig. 5). **b,** Fitted kobs for Rab7 as a function of the ceTBC1D5 concentration at pH 7.0 *in vitro*. Solid lines are linear fits to kobs as a function of ceTBC1D5 concentration. **c,** Catalytic efficiency (kcat/KM) of purified ceTBC1D5 on Rab7 increased over the pH range and with a profile consistent with histidine deprotonation and remained constant after pH 7.5(n = 2-5 experiments). The solid line is a fitted model function for the pH dependence of one or more titratable groups with a common pKa (see Methods). The mean for the fitted pKa value is 5.95 ± 0.02. **d,** AlphaFold 3^65^ model shows a TBC1D5 dimer (upper top, green and turquoise) directly binds to the C-terminus of a NHE6 dimer (cyan and purple), which is embedded in a lipid bilayer. GTP-Rab7 (grey) is correctly positioned on the catalytic domain of TBC1D5 in this model. Note location of an interfacial histidine proximal to the invariant Arginine finger/aspartate and Glutamine finger/tyrosine. (see Extended Data Fig. 6e)**e,** Alignment of representative TBC domain orthologs and paralogs from the three major phylogenetic supergroups^81^. **f,** The co-localization of NHE6 (cyan), TBC1D5 (magenta), and Rab7 (yellow) in primary neurons. Triple labelling is shown in white. An orange box indicates the inset. On average, 33.74 % (± 4.09 %) of Rab7-punctae were co-stained with TBC1D5 and NHE6. Means from each animal (big dots, 3 WT pups) overlay the entire dataset (small dots, n = 28 neurons from three independent experiments). Data are represented as mean ± SEM. Scale bar, 5 μm. Scale bar for the inset, 1 μm.

Based on our observations, we used AlphaFold 3 to predict a structural model for a dimeric NHE6-TBC1D5-Rab7-GTP complex with 2:2:2:2 stoichiometry (Fig. 5d)^65^. In this model, the TBC domain of TBC1D5 dimer binds to the C-terminal region of the NHE6 dimer, which is embedded in a lipid bilayer. The TBC1D5-Rab7-GTP interface is consistent with previous crystallographic and biochemical studies of the Gyp1-Rab33 and TBC1D20-Rab1 complexes, including the dual catalytic Arginine and Glutamine fingers of the TBC domain with respect to the hydrolytic site^66^. Within this interface, there is a conserved histidine (His 207) located proximal to these Arginine and Glutamine fingers as well as other invariant hydrolytic site residues. This interfacial histidine (His 207) is conserved in distant TBC1D5 and TBC1D20 orthologues but not in TBC domains generally (Fig. 5e), suggesting a putative role as a pH-sensing amino acid to regulate pH-dependent Rab7 GAP.

We also performed triple-immunofluorescence staining to investigate NHE6- TBC1D5-Rab7 co-localization in neurons using high-resolution microscopy with the NSPARC detector (Fig. 5f). We stained primary WT neurons with anti-Rab7, TBC1D5, and NHE6 antibodies. Consistent with our AlphaFold 3 model, we observe substantial co-localization of all three proteins together on endosomes in primary neurons. On average, we found 33.74 % (± 4.09 %) of Rab7-positive late endosomes were co-stained with TBC1D5 and NHE6 in primary neurons. In summary, our data support the conclusion that TBC1D5 serves as a pH sensor that detects changes in local pH mediated by proton efflux via NHE6, thereby regulating Rab7 activity through a direct mechanism involving robust inactivation of Rab7 GAP activity with decreasing pH.

### Knock-down of the Rab7 GAP TBC1D5 rescues endosomal phenotypes in NHE6- null neurons

To determine the extent to which increased TBC1D5 activity on Rab7 causes disrupted Rab7 function in late endosome maturation and trafficking in NHE6-null neurons, we established a lentiviral system for acute knock-down of TBC1D5 levels. We transduced TBC1D5 shRNA into primary WT and NHE6-null neurons at 2 days *in vitro* (DIV) and confirmed the decreased expression of TBC1D5 protein by western blot (Fig. 6a) and immunostaining with an anti-TBC1D5 antibody (Extended Data Fig. 6). We then monitored the movement of mEmerald-Rab7 endosomes in primary WT and NHE6-null neurons transduced either with scrambled shRNA or TBC1D5 shRNA (Fig. 6b). NHE6- null neurons transduced with scrambled shRNA displayed fewer retrograde Rab7 endosomes and a greater number of stationary Rab7 endosomes compared to WT neurons, as we observed previously in non-transduced neurons (Fig. 1d). However, transduction of TBC1D5 shRNA lentivirus in NHE6-null neurons reversed these phenotypes. Specifically, transduction with TBC1D5 shRNA in NHE6-null neurons significantly increased the number of retrograde Rab7 endosomes while decreasing the number of stationary Rab7 endosomes compared to NHE6-null neurons transduced with scrambled shRNA. We also performed the FRAP assay in primary WT and NHE6- null neurons transduced with scrambled or TBC1D5 shRNA (Fig. 6c). The recovery of Rab7 was significantly increased in primary NHE6-null neurons transduced with TBC1D5 shRNA, compared to primary NHE6-null neurons transduced with scrambled shRNA. Lastly, we had previously shown that endosome-lysosome fusion, a key step in endosome maturation, was impaired in NHE6-null neurons^21^. Furthermore, TBC1D5 plays a key role in endosomal fusion with lysosomes^53^. Late endosome fusion with lysosomes was monitored in TBC1D5 shRNA transduced neurons (Fig. 6d). The overlap of endosomes with lysosomes, signifying endosome-lysosome fusion, was significantly increased in primary NHE6-null neurons transduced with TBC1D5 shRNA, as compared to NHE6-null neurons with scrambled shRNA. Overall, these data indicate knock-down of TBC1D5 rescues Rab7 functions in NHE6-null neurons, demonstrating that overactive TBC1D5 and dysregulated Rab7 are linked to endosome and trafficking defects in the absence of NHE6 function.

**Fig. 6:**
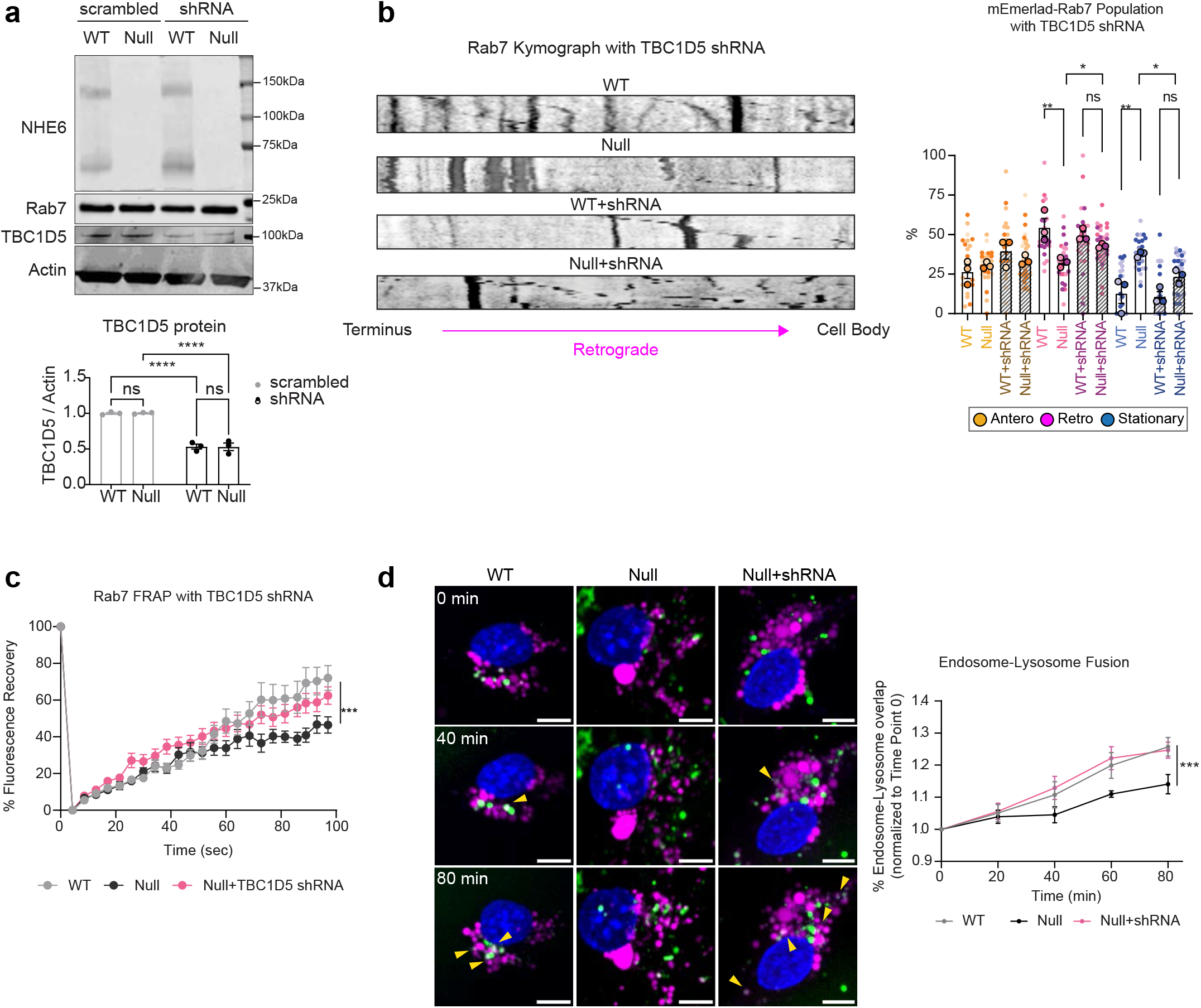
Knock-down of the Rab7 GAP TBC1D5 rescues endosomal phenotypes in NHE6-null rat neurons. **a,** Reduced protein expression of TBC1D5 in primary littermate WT and NHE6-null neurons, with lentiviral transductions of anti-TBC1D5 shRNA, as compared to scrambled shRNA control transductions The expression of NHE6 and Rab7 was not affected by the shRNA transduction (three independent experiments). Two-way ANOVA with Fisher’s LSD test. **b,** Representative kymographs from primary littermate WT and NHE6-null neurons with scrambled or TBC1D5 shRNA show the movement of mEmerald-Rab7 endosomes. Retrograde direction is indicated. Scale bar, 1 μm. The number of retrograde Rab7 endosomes in primary NHE6-null neurons transduced with TBC1D5 shRNA was increased compared to NHE6-null neurons transduced with scrambled shRNA. Also, the number of stationary Rab7 endosomes was decreased in primary NHE6-null neurons transduced with TBC1D5 shRNA. Means from each animal (big dots, WT = 3, Null = 3 pups) are overlay the entire dataset (small dots, n = 21-31 neurons from three independent experiments) and used for statistical analysis. Animals from same litters are color-coded. Ordinary one-way ANOVA with Tukey’s HSD. **c,** The recovery rate of mEmerald-Rab7 in primary NHE6-null neurons transduced with TBC1D5 shRNA exhibited significant increase relative to WT neurons. The recovery rate of mEmerald-Rab7 in NHE6-null neurons transduced with TBC1D5 shRNA was comparable to WT neurons or NHE6-null neurons with scrambled shRNA (6 pups in total for each genotype, n = 20-35 neurons from 3 experiments). **d,** The increased endosomes-lysosomes fusion in primary NHE6-null neurons transduced with TBC1D5 shRNA. Shown are still images at different time points from live imaging of the endosome-lysosome fusion assay on primary littermate WT and NHE6-null neurons transduced with scrambled or TBC1D5 shRNA at different time points. The level of endosome-lysosome fusion is presented as the percentage fold change in overlap between the lysosome label (magenta) and the endosome label (green), from time point 0 for the same animal. Yellow arrows indicate the fusion events. (WT = 5, Null = 5 pups, n = 10,000-14,000 neurons imaged from each pup) Linear mixed model was conducted. Scale bar, 5 μm. Data are represented as mean ± SEM. ****p < 0.0001, ***p < 0.001, **p < 0.01, and *p < 0.05.

## Discussion

Two critical processes during endosomal maturation include the progressive acidification of the endosomal lumen and the regulation of Rab GTPase function on the cytoplasmic side. The linkage of these two mechanisms in governing endosome maturation has not been previously demonstrated. Furthermore, “inside-out” endosome signaling mechanisms - that is communication to the cytoplasmic side reporting the process of intra-endosome acidification - have been hypothesized and long sought-after but have remained elusive^1,67^. In this study, we provide abundant evidence that proton efflux, mediated by a Na^+^/H^+^ Exchanger, acts as an “inside-out” signal heralding endosome acidification. We propose that this process uses the passive proton gradient extending from the proton efflux site to regulate a pH-dependent Rab GAP to synchronize critical lumen and cytoplasmic processes required for endosome maturation. Thereby, we identify a new function of endosomal NHEs in this signaling mechanism. Furthermore, our studies link the classically defined mechanisms of lumen acidification with Rab GTPase cycling during endosome maturation. This mechanism likely functions down downstream of Rab conversion^17,18,19^, potentially as a brake on Rab7-GTP accumulation until lumen acidification reaches the optimal pH range. Figure S7 provides a schematic model for the mechanism studied here wherein proton efflux through NHE6 regulates Rab7 function causing endosome maturation by inactivating the Rab7 GAP TBC1D5.

In order to study proton efflux through NHE6, we engineered a new mouse model (NHE6-ED) harboring two-point mutations that inactive the exchanger domain of NHE6 ^27,39^. Importantly, these mutations impair proton efflux, as we demonstrate that NHE6- ED intra-endosomal pH is over-acidified, akin to NHE6-null neurons. Using this NHE6- ED model, we find that compromised NHE6 proton efflux disrupts the trafficking and GTP-GDP cycling of Rab7-positive endosomes. This result also mimics our observations in NHE6-null neurons. In our studies, defects in Rab7 GTPase cycling in the context of NHE6 mutations are particularly convincing given that we observe these defects with two distinct NHE6 mutations, using multiple complementary methods, in two rodent species. Furthermore, the interaction between NHE6 and TBC1D5 is well-supported by reciprocal co-immunoprecipitation, as well as co-localization studies using super-resolution microscopy. In our triple-labeling studies, we observe a substantial fraction of triple-localization of NHE6, Rab7, and TBC1D5. A prior study in yeast that shows interaction between Nhx1 (the NHE6 homologue) and Gyp6 (the TBC1D5 orthologue) provides additional support for the NHE6 and TBC1D5 interaction^42,59^.

Knock-down of TBC1D5 restores Rab7 endosomes trafficking, the GTP-GDP dynamics of Rab7, and endosome-lysosome fusion^21^. The knock-down experiment provides important functional evidence for our proposed mechanism.

Our biochemical studies of TBC1D5 indicate that the Rab GAP activity is regulated by changes in pH. We observe that the catalytic activity rests near its maximum at cytosolic pH; however, falls sharply as pH is decreased, with a profile that is consistent with histidine protonation. Indeed, the observed pKa is nearly identical to that of histidine. The TBC domain of TBC1D5 contains 19 histidines, four of which are conserved in the worm orthologue RBG-3. Interestingly, one of these conserved histidines is located proximal to the catalytic Arginine and Glutamine fingers ^66^ as well as other invariant hydrolytic site residues in an AlphaFold 3 ^65^ structural model of a NHE6-TBC1D5-Rab7-GTP dimer. This interfacial histidine is conserved in distant TBC1D5 and TBC1D20 orthologues, but not in TBC domains generally. Notably, the corresponding histidine occupies an equivalent location in the crystal structure of the TBC1D20-Rab1 complex^66^. The interfacial histidine has no known function, and consequently, conservation in distant orthologues is intriguing but at present lacks explanation. A plausible possibility is that protonation of the interfacial histidine directly impairs catalytic activity by introducing an energetically unfavorable, uncompensated positive charge, incompatible with the buried location of the interfacial histidine in the complex with Rab7. Our immunostaining studies support the presence of a NHE6- TBC1D5-Rab7 complex on endosomes in primary neurons. The AlphaFold 3 model further predicts that a dyad symmetric TBC1D5 dimer would be positioned in close proximity to the cytoplasmic surface of the dyad symmetric NHE6 dimer, consistent the proposed role of TBC1D5 as a sensor of proton efflux through the NHE6. Additional studies are required to determine whether the interfacial histidine is directly responsible for the observed pH dependence of the catalytic activity or whether histidine residues outside the interface with Rab7 contribute to and/or mediate the effect through an allosteric mechanism.

Loss-of-function mutations in NHE6 causes an X-linked neurological disorder Christianson syndrome (CS). CS patients display developmental delay, postnatal microcephaly, absent speech, progressive ataxia and epilepsy along with neurodegenerative disease^32,68,69^. There is prominent axonal pathology in both human postmortem and animal models^34,70^. Disrupted trafficking, and accumulation of enlarged and as well as clusters of endolysosomes have been implicated in the pathogenesis of other neurodegenerative disorders including Alzheimer’s disease^24,44,48,71^. In our NHE6- null and NHE6-ED neurons, we observed disrupted trafficking and the accumulation of enlarged Rab7-positive endosomes. These Rab7 defects may cause “traffic jams” along the axons and dendrites, leading to cascading downstream effects during aging and neurodegeneration^55,72^. In previous studies, we have shown that NHE6-null rat brains exhibited the accumulation of enlarged endolysosomes, preceding autophagosomes dysfunction^34^. In that study, we also observed that gliosis, axonal loss, neuronal loss and tau deposition occur in NHE6-null rat brains.

Our model for the regulation of TBC1D5 by NHE6-mediated proton efflux involves dynamic fluctuations in pH-microdomains (Extended Data Fig. 7). One primary limitation of this current work is the inability to measure the pH of the cytoplasmic microenvironment adjacent to late endosomes. Proton efflux from endosomes and other organelles rapidly dissipates due to the buffering capacity of the cytosol^73^. Currently, there are no pH indicators available to measure this rapid proton change between endosomes and the cytosol at the required spatial and time resolution in cells.

Additionally, future studies may seek to expand on this molecular mechanism by identifying other Rab GAPs or guanine nucleotide exchange factors (GEFs) that may be pH dependent. Furthermore, study of other endosomal NHEs in proton-mediated signaling mechanisms, and identification of other pH-dependent targets, may expand on this conceptually new mechanism in endosome trafficking.

## Methods

### Rats and mice

All animal work was conducted under the guidelines of the Center for Animal Resources and Education (CARE) with a protocol (IACUC 21-09-0012), approved by the Brown University Institutional Animal Care and Use Committee (IACUC). All experimental procedures were consistent with the US National Institutes of Health Guide for the Care and Use of Laboratory Animals (National Research Council, 8th edition). Animals were kept at a 12/12-hour light/dark cycle with ad libitum access to food and water. As NHE6 is located on the X chromosome, only male rats were used for this study. Heterozygous females were crossed with wild-type males for breeding. Tails from rats and mice were clipped and externally genotyped (Transnetyx, Cordova TN).

NHE6-null rats were originally described in our previous study^34^. Age of animals used for this study were indicated in each figure and/or figure legends. A mouse model of human E255Q/D260N variant (mouse E235Q/D240D) was generated for this study. This line is referred as NHE6 efflux-defective (NHE6-ED) mutant line. Mouse lines were generated using CRISPR/Cas9 genome editing (Mouse Transgenic and Gene Targeting Facility of Brown University) and are described for the first time herein. We used a guide RNA (5’- GGAAAGTGTCCTCAATGACG-3’) to generate both mutations. Mouse lines were generated on the C57BL/6J mouse background. One male founder and two female founders were generated and used for breeding. Studies were conducted in offspring from each of the different founders. Targeting of constructs and presence of mutations were confirmed by PCR genotyping and Sanger sequencing (Extended Data Fig. 4C). The following primers were used for PCR genotyping: forward primer (5’- AGCTGTGGAGGGATATGTGC-3’) and reverse primer (5’- TCAGAGCAGGGCAGAAAGAC-3’). The expression of protein was confirmed by western blot (Extended Data. 4d).

### Tissue preparation for staining and protein assays

For immunofluorescence staining, animals were anesthetized using pentobarbital (100 mg/kg). Animals were transcardially perfused using phosphate-buffered saline (PBS) followed by 3.7 % formaldehyde (Sigma-Aldrich). Following cardiac perfusion, the brain was stored in 3.7 % formaldehyde fixative at 4 °C overnight. For immunofluorescence, the brain was transferred to 30 % sucrose solution at 4 °C and stored until sinking.

Frozen brain samples were coronally sectioned at 50 μm thickness on a sliding microtome (Thermo Scientific #HM430) at -20 °C. Sections were serially collected and stored in 24-well plates containing a cryoprotectant solution (30 % sucrose, 30 % ethylene glycol, 1 % Polyvinyl-pyrrolidone in PBS) at -80 °C until use.

For protein extraction, wild-type and NHE6-null rats were euthanized with CO2 and brains were removed, dissected and cut in half. Each hemisphere was weighed, snap frozen and homogenized for protein extraction in NP-40 lysis buffer (Thermo Scientific # J60766.AP).

### Stereological sectioning and analysis

Anatomical brain regions of interest (ROI) were confirmed according to the rat brain atlas^74^. Three to five sequential coronal sections 300 μm apart between bregma -1.88 mm to -6.72 mm were taken per each animal. All procedures including staining, image acquisition, and analysis were performed blind to genotype and age. Only male animals were used.

For counting the number and size of SMI32 or Rab7, five serial coronal sections 300 μm apart from WT and NHE6-null rats were taken for staining and analysis. Images for each section were randomly taken within the defined ROI, i.e., CC and CA1 using 63x (NA 1.40) oil immersion objective (Olympus). The number and size of SMI32 or Rab7 per each image was manually counted by an investigator using Fiji ImageJ who was blinded to genotype and age of animals. If SMI32 or Rab7 puncta’s diameter size is larger than 1.5 μm, we defined them as swollen/enlarged Rab7. The number of swollen SMI32 puncta was divided by the total puncta number to present the percentage (%).

For counting the Rab7 presence in the Tau-stained axons, we generated a mask from Rab7 staining and overlaid the Rab7 mask to tau staining in Fiji ImageJ. The number of Rab7 and % of Rab7 covered area in this mask were only counted.

All tissue sections were imaged under a confocal laser scanning microscope (Olympus FV3000) with the same imaging settings between control and mutant animals. Mutant and control images were equally adjusted in Fiji ImageJ software for brightness and contrast to ensure accurate data analysis. All data analysis was performed by someone blind to the genotypes and age of the animals using Fiji ImageJ.

### Primary neuronal culture

Hippocampi were dissected from P0-P1 rats or mice dissociated with papain dissociation kit (Worthington #LK003150) as previously described^27^. Only male rats and mice and their wild-type littermate controls were cultured for this study. Briefly, the brains were removed from the skull, and hippocampi were dissected on ice in prechilled Hank’s balanced salt solution (HBSS; Invitrogen #14025076). All dishes were pre-coated with 0.1 mg/mL of Poly-D-lysine (PDL; Sigma #A3890401) for overnight at 37 °C. Cells were cultured in neurobasal media (Invitrogen #21103049) supplemented with 2 % B27 (Invitrogen #17504044), 1 % glutamax (Invitrogen # 35050061), and penicillin/streptomycin. Cells were seeded at densities of 28,000 cells per well in the 96- well plate (Perkin Elmer #6055302) for high-throughput imaging. 120,000 cells per dish were seeded for #1.5 14 mm glass bottom dishes (Cellvis #D35-14-1.5-N) for live-imaging. For immunofluorescence, 40,000 cells were seeded on each #1.5 12 mm PDL-coated German glass cover slips (neuVitro #GG-12-1.5-PDL) and placed in 24-well plates. 1.2x10^6^ cells per well were seeded for 6-well plate for western blotting. For shRNA lentivirus transduction, scrambled shRNA and TBC1D5 shRNA (MOI = 2) were transduced in primary neurons at DIV 2 and incubated for 72 hours prior to protein extraction.

### Live-imaging and fluorescence recovery after photobleaching (FRAP)

Primary neurons from male controls and mutants were grown on PDL-coated dishes. Seeding density are indicated above. Cells were transfected with mEmerald-Rab7 a day prior to imaging at DIV 7-9. Cells were incubated with 1 μg of plasmid DNA mixed with 2 μL of lipofectamine 2000 (Invitrogen #11668019) on 37 °C for 2 hours. After the incubation, the transfection medium was replaced with the culture medium pre-saved from the cultures prior to the transfection. On a day of imaging, primary neurons were incubated in imaging media containing phenol red-free neurobasal media (Thermo Fisher # 12348017) supplemented with 2 % B27, 1 % glutamax, and penicillin/streptomycin. mEmerald-Rab7a-7 was a gift from Michael Davidson (Addgene plasmid # 54244; http://n2t.net/addgene:54244 ; RRID:Addgene_54244). TBC1D5 shRNA lentivirus (GeneCopoeia #LPP-RSH087343-LVRH1MP-200) was transduced at DIV 2 (MOI = 2) for 72 hours. For live-imaging, images were acquired on Olympus 3000 confocal microscope using 63x (NA 1.40) oil immersion objective (Olympus) with 2x digital zoom-in. Live cells were imaged in a temperature-controlled chamber (37 °C) at 5 % CO2 at 1 frame per second for 90 seconds. The total 4 μm of Z-stack was acquired in 0.44 μm step size. Only the neurons with identified axons were imaged. Images were deconvoluted and maximum projected in Olympus Cell Sense software. Kymograph were acquired and analyzed with “KymoAnalyzer” ^75^ Plugin in Fiji ImageJ. For FRAP, region of interest (ROI) was pre-determined in the neuronal cell bodies with stationary endosomes. 3 seconds of pre-bleached images were acquired. Photobleaching was performed with 5 % of 488 nm laser for 500 milliseconds with no interval delay. After bleaching, images were live-recorded for 100 seconds at 1 frame per second. Deconvoluted images were analyzed using “Stowers” Plugins in Fiji ImageJ.

### RILP assay

Brains from controls and mutants at 6-8 weeks were lysed. 500 μg of protein lysates were incubated with 5 μg of His-RILP (NovoPro Labs #512098) or His-Myc using His Pull-down kit (Thermo Scientific #21277) for 2 hours at 4 °C. Samples were then eluted with 300 mM imidazole and mixed with loading buffer and reducing agent before they were boiled at 95 °C for 5 minutes. Western blotting was performed to detect Rab7 from the elutants.

### GTP-Rab7 pull-down assay

This assay was adopted and modified from the Rab5-GTP pull-down assay^55^. Brain lysates from WT and NHE6-null rats at 6-8 weeks were centrifuged at 13,000 xg for 10 minutes at 4 °C. The supernatant was transferred to a new tube and the protein concentration was measured by BCA assay. 200 μg of protein lysates was incubated with 50 μL of GTP-agarose (Sigma #G9768) for 2 hours at 4 °C. The samples were spun down at 5,000 xg at 4 °C for 1 minute. The supernatant was discarded, and the pellet was washed with lysis buffer for 3 times. After the last wash, the samples were mixed with loading buffer and reducing agent and boiled at 95 °C for 5 minutes. Western blotting was performed to detect Rab7.

### Immunofluorescence

Primary neurons from controls and mutants were grown on PDL-coated German glass cover slip #1.5. Seeding density are indicated above. Primary neurons were washed three times with ice-chilled PBS and fixed in 3.7 % paraformaldehyde for 10 minutes. Cells were again washed with PBS for 5 minutes, three times, and permeabilized with PBS containing 0.1 % Triton-X100 (PBS-T; Sigma) for 10 minutes. Cells were blocked in blocking buffer (PBS-T containing 10 % goat serum) for one hour at room temperature. All primary antibodies except Rab7 antibodies were diluted in the blocking buffer and incubated for overnight, 4 °C. Rab7 antibodies were incubated for more than 48 hours at 4 °C. Secondary antibodies and Hoechst 33342 were diluted in the blocking buffer as well and incubated for one hour at room temperature. Cells were washed three times with PBS for 5 minutes for each wash and mounted with ProLong Gold Glass Antifade mountant (Invitrogen #P36984). Images were acquired on Nikon AX R with NSPARC detector using 63x (NA 1.40) oil immersion objective with 4x digital zoom-in or CellCarrier-96 Ultra microplates (PerkinElmer). The quantification was performed in Opera Phenix Harmony (PerkinElmer) to calculate the Mander’s coefficient.

### Immunoprecipitation and western blotting

For immunoprecipitation and western blots, whole brains from WT and NHE6-null rats were lysed in ice-chilled RIPA buffer (50 mM Tris pH 7.4, 150 mM NaCl, 1 mM EDTA, 10 % of TritonX-100, 0.1 % of sodium deoxycholate, 0.1% of sodium dodecyl sulfate) with complete proteinase inhibitor cocktail (PIC; Sigma), 10 μL/mL phenylmethylsulfonyl fluoride (PMSF; Sigma), and phosSTOP phosphatase inhibitor (Sigma #4906837001). Lysates were incubated on ice for 30 minutes and then spun down at 13,200 xg, 4 °C for 15 minutes. For immunoprecipitation, 200 μg of samples were pre-cleared for 1hr on end-over-end rotator with 10 μg of M-270 Epoxy Dynabeads (Life Technologies #14302D) at 4 °C. The supernatants were incubated with 10 μg of NHE6 antibody-conjugated Dynabeads on the rotator at 4 °C for overnight. After the incubation, the samples were washed three times with ice-chilled lysis buffer. Samples were mixed and boiled at 95 °C with NuPage 4× LDS sample buffer (Invitrogen # NP0007) and NuPage 10x sample reducing agent (Invitrogen # NP0009). The samples were run on NuPage 4- 12 % SDS-PAGE gel (Invitrogen # NP0321BOX) using MOPS buffer and transferred to nitrocellulose membranes. The primary antibodies: Rabbit anti-NHE6 (1:1000, Covance), rabbit anti-TBC1D5 (1:1000, Novus Biological # NBP1-93653), and mouse anti-actin (1:5000, Proteintech) were used. After three washes with TBS containing Tween20 (TBS-T), IRDye 680W and 800W goat anti-rabbit and anti-mouse secondary antibodies were incubated with samples for 1 hour at room temperature. All the membranes were visualized with the LiCor Odyssey Clx Infrared Imaging System.

### Structure illumination microscopy (SIM)

Primary neurons from NHE6-null rats and their wild-type littermates were grown on # 1.5 PDL-coated coverslips and fixed in 3.7 % paraformaldehyde for 10 minutes at room temperatures. The same immunofluorescence staining procedures were followed as described above. Cells was imaged on 63x oil immersion oil (NA 1.46) using the DeltaVision OMX SR imaging system (GE).

### Endosomal pH measurement

Primary neurons were seeded at densities of 28,000 cells per well on CellCarrier-96 Ultra microplate (PerkinElmer). The assay was conducted as previously described in detail^63^. Briefly, primary neurons at DIV 7 were first starved in EBSS at 37 °C for 30 minutes. EBSS was then replaced with Neurobasal A media with 33 μg/ml fluorescein isothiocyanate (FITC)-conjugated transferrin (Thermo Fisher Scientific #T2871), which is pH sensitive, and 33 μg/ml Alexa Fluor 546-conjugated transferrin (Thermo Fisher Scientific # 23364), which is pH insensitive. After 10 minute-incubation, the cells were washed twice with warm PBS, placed in phenol red-free Neurobasal media, and imaged live under 60x water objective using an Opera Phenix High-Content Screening System. To generate the standard curve for use in determining endosomal pH, similar procedures were followed; however, neurons were imaged in standard buffer solutions as described previously^63^.

### Endosome-lysosome fusion assay

Primary neurons were seeded at densities of 28,000 cells per well on CellCarrier-96 Ultra microplates (PerkinElmer). The assay was conducted as previously described in detail^21,76^.

### Measure the catalytic activity of TBC1D5

*Constructs.* Constructs corresponding to residues 5-176 of mouse Rab7 (NM_009005) and 15-449 of worm TBC1D5 (NM_065578; also known as RBG-3) were amplified and inserted into BamHI/SalI sites of pET15 modified to incorporate an N-terminal 6xHis tag (MGHHHHHHGS).

*Expression and Purification.* Constructs transformed into BL21(DE3)-RIPL cells (Novagen) were expressed in 2xYT media supplemented with 100 mg/L ampicillin. After growth at 37 °C to OD600 ∼0.2, the temperature was lowered to 22 °C until OD600 ∼0.4. Expression was induced with 0.05 mM IPTG for 16 hours. Cell pellets were resuspended in 50 mM Tris, pH 8.0, 150 mM NaCl, 2 mM MgCl2, 0.1 % 2- mercaptoethanol, 0.1 mM PMSF, 0.01 mg/mL protease free DNAse I (Worthington), and 0.2 mg/mL lysozyme. After incubation on ice for 1 hour followed by sonication, the lysates were supplemented with 0.5 % Triton X-100 and centrifuged at 28,000 ×g for 1 hour at 4 °C. After purification over NiNTA-sepharose HP, HiTrap QHP and Superdex 200 columns, proteins were estimated to be >95 % pure by SDS-PAGE.

*Nucleotide Loading.* Rab7 (1 mg/mL) was incubated for 2 hours at room temperature with a 20-fold molar excess of GTP in 20 mM Tris, pH 7.5, 100 mM NaCl, 5 mM EDTA, and 1 mM DTT. Excess nucleotide was removed using a 10 mL D-Salt column (Thermo-Fisher) equilibrated with 20 mM Tris, pH 7.5 and 150 mM NaCl.

*GAP Assays.* Single turnover GAP assays were performed as described^77^, using the increase in intrinsic tryptophan fluorescence of Rab7 to monitor the rapid conformational switch from the GTP- to GDP-bound state resulting from GTP hydrolysis^16^. Reactions were initiated by mixing 50 μL of 4 μM Rab7-GTP in 20 mM buffer, 150 mM NaCl with an equal volume of the same buffer containing 10 mM MgCl2 and 0 - 2.56 μM RBG-3.

Buffers were sodium acetate (pH 4.5-5.5), PIPES (pH 6.0-7.0), and Tris (pH 7.5-8.5). Dispensing, mixing and dilutions were performed with a Precision 2000 automated multichannel pipetting system (Biotek). The increase in Rab7 intrinsic tryptophan fluorescence accompanying GTP hydrolysis was continuously monitored in half area microplates (Corning) using a Tecan Spark microplate spectrometer with excitation at 290 nM, emission at 340 nm, and a bandwidth of 10 nM. Fluorescence time courses Ft were corrected for a small amount of photobleaching and fit with the exponential model function

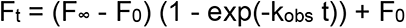

where kobs is the observed rate constant and F0 and F∞ are the fluorescence intensities at t = 0 and t→∞, respectively. A linear term was included to account for a small increase in fluorescence observed at higher GAP concentrations. Catalytic efficiencies (kcat/KM) were obtained from the slopes of linear fits to kobs vs. t. The pH dependence of the catalytic efficiency was fit with the model function:

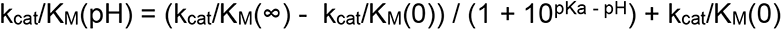

where kcat/KM(pH) is the observed rate constant as a function of pH and kcat/KM(0) and kcat/KM(∞) are the fluorescence intensities at pH → 0 and pH → ∞, respectively. This model describes the pH dependence for one or more titrating groups with a common pKa such that kcat/KM(0) corresponds to the catalytic efficiency of the fully protonated state and kcat/KM(∞) to that of the fully unprotonated state. For fitting, kcat/KM and kcat/KM(∞) were treated as adjustable parameters whereas kcat/KM(0) was fixed at zero since fitting it resulted in a small, nonphysical negative value. The Mac OSX application DELA was used for data processing, analysis and relevant figures^77,78^.

### AlphaFold 3 structural modeling

The AlphaFold 3 structural model for the dimeric NHE6-TBC1D5-Rab7 complex was generated using the AlphaFold 3 server^65^, with two copies of the amino acid sequences for human NHE6 (NP_001036002), human TBC1D5 (NP_001127853), and human Rab7 (NP_004628), two copies of GTP, and two copies of Mg^2+^. Figures for the model were rendered with PyMOL.

### Electron microscopy (EM)

Genotypes and conditions of animals were blinded during image collection and analyses. Tissue was collected and processed for standard electron microscopic analysis as previously described^79^.

### Quantification and Statistical Analysis

All statistical methods and number of biological replicates used for this study are described in the figure legends and the Methods. In this study, values from each biological replicate are clustered in different color codes and plotted as a small dot. Means from each biological replicate are overlaid on the top of the full dataset as a bigger dot. All the statistical analyses were calculated across biological replicates (represented as bigger dots), not the entire dataset (represented as small dots)^80^. Data are represented as means ± SEM throughout the manuscript. Asterisks represent p-value as follows: **** < 0.0001, *** < 0.001, ** < 0.01, and * < 0.05.

## Acknowledgements

This research was supported by funding for YL (K99AG076868) and EMM (R01NS121618, R01AG087455, R01NS113141). We thank Dr. Diane Barber at University of California San Francisco, and Dr. Nicholas Fawzi at Brown University for feedback and insightful discussion.

## Author information

YL designed, conducted, and analyzed experiments. YL also wrote the manuscript. EMM designed and supervised experiments and wrote the manuscript. HR, LM, QO, MF, MS, JD, and DGL designed, conducted, and analyzed experiments.

## Ethics declarations

The authors declare no competing interests.

## Materials & Correspondence

Further information and requests for resources and reagents should be directed to and will be fulfilled by Eric Morrow.

---

**Extended Data Fig. 1.**
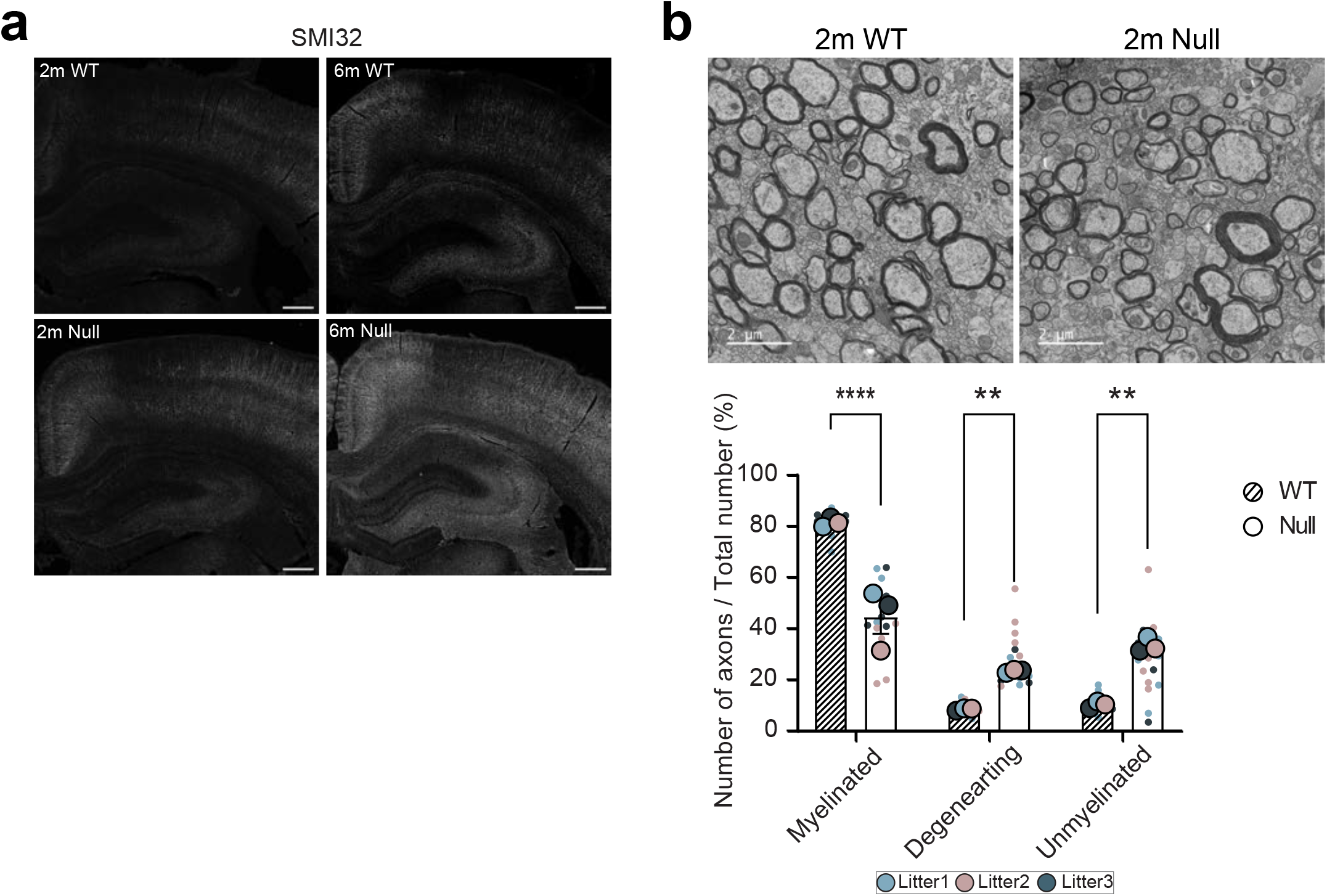
Axonal degeneration in NHE6-null rat brains. **a,** Representative images of the brain sections from littermate WT and NHE6-null rat brains at 2 and 6 months. Brain sections were stained with SMI32 (axonal damage marker, gray). **b,** Representative electron microscopy images of the corpus callosum from littermate WT and NHE6-null rat brains at 6-7 weeks. The total number of axons, and the number of myelinated, demyelinated, and degenerating axons for each image were counted (3 animals per each genotype). Scale bar, 2 μm. Two-way ANOVA with Tukey’s HSD was performed. Data are represented as mean ± SEM. ****p < 0.0001, ***p < 0.001, **p < 0.01, and *p < 0.05.

**Extended Data Fig. 2.**
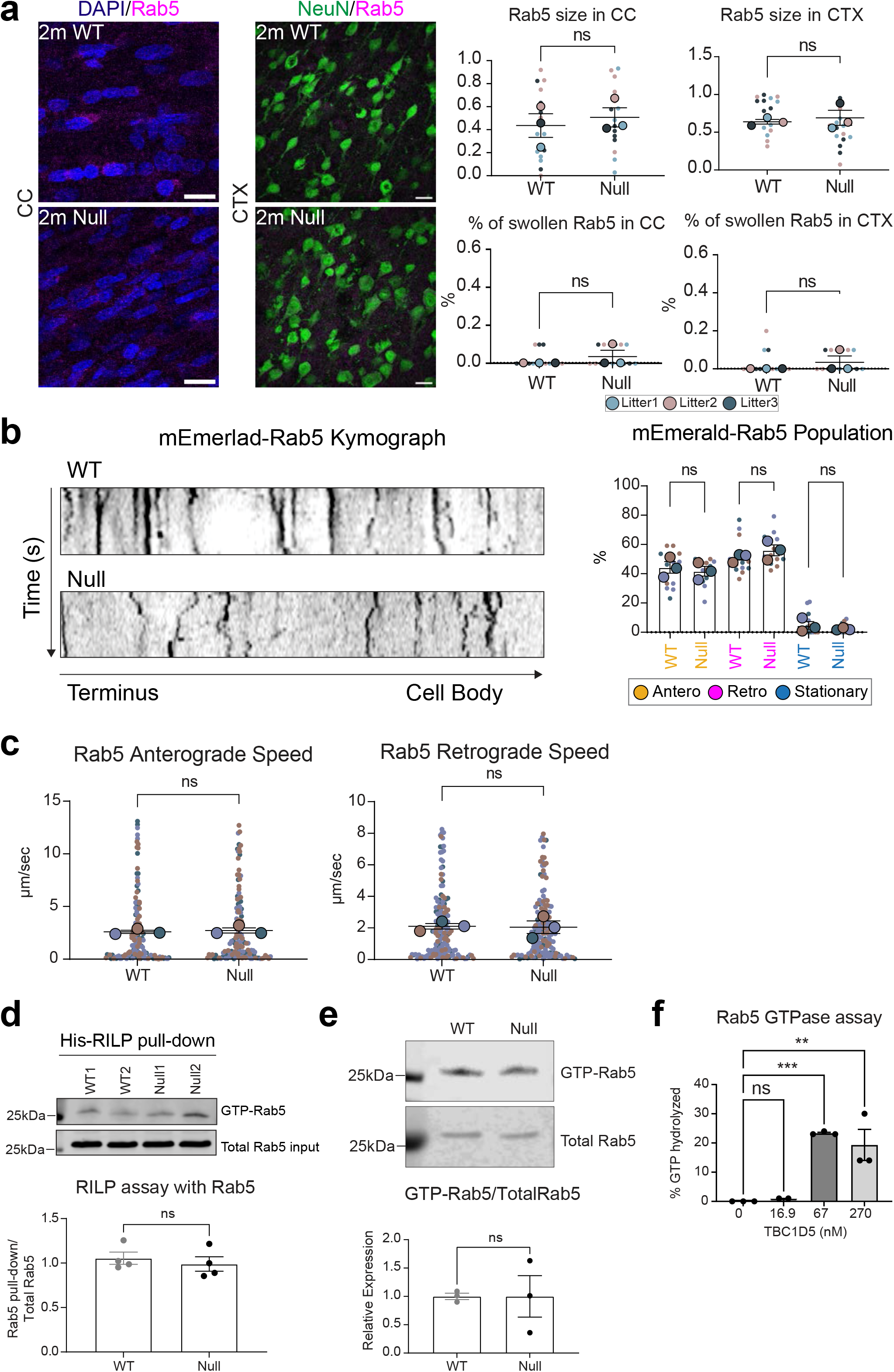
Early endosome defects were not detected in NHE6-null rats. a, Representative images from the corpus callosum and cortex sections of littermate WT and NHE6-null rat brains at 2 months. Sections were stained with Rab5 (early endosomal marker, magenta). NeuN was used as a neuronal marker (green). No differences in size and number of enlarged Rab5 were detected between WT and NHE6-null rat brains. The number of Rab5 was counted and divided by the total number of Rab5 puncta (%). Also, the average size of Rab5 puncta was measured. Means from each independent experiment (big dots, WT = 3, Null = 3 animals) overlay the entire dataset (small dots, 5 sections per each animal) and used for statistical analysis. Animals from same litters are color-coded. Two-tailed unpaired t-test with Welch’s correction was performed. Scale bar, 10 μm. **b,** Representative snapshots and kymographs from primary littermate WT and NHE6- null neurons show the movement of mEmerald-Rab5. Retrograde direction is from left (terminus) to right (cell body). Scale bar, 1 μm. There were no significant differences in the number of anterograde, retrograde, and stationary Rab5 endosomes in primary WT and NHE6-null neurons. Means from each animal (big dots, WT = 3, Null = 3 pups) overlay the entire dataset (small dots, n = 18-20 neurons from four independent experiments) and used for statistical analysis. Animals from same litters are color-coded. Ordinary one-way ANOVA with Tukey’s HSD was performed. **c,** Speed of anterograde and retrograde mEmerald-Rab5 endosomes in primary WT and NHE6-null neurons. Stationary Rab5 endosomes were not included for the calculation. There were no differences in speed. **d,** RILP pull-down assay with Rab5. There were no differences in the amount of RILP- bound Rab5 (GTP-Rab5 pull-down) between WT and NHE6-null rat brains at 2 months. Lysates from WT and NHE6-null rat brains were incubated with His-RILP recombinant proteins for the pull-down assay. Once samples were eluted, Rab5 was detected by western blot. The densitometry of GTP-Rab5 was divided by the total Rab5 for the quantification (4 rat brains for each genotype). Two-tailed unpaired t-test with Welch’s correction was performed. **e,** GTP agarose assay with Rab5. There were no differences in the amount of GTP-bound Rab5 between littermate WT and NHE6-null rat brains at 2 months. Lysates from WT and NHE6-null rat brains were incubated with GTP agarose to enrich GTP-bound protein fractions. Rab5 was detected in the GTP-enriched fraction by western blot. The densitometry of GTP-bound Rab5 was divided by the total Rab7 for the quantification (3 rat brains for each genotype). Two-tailed unpaired t-test with Welch’s correction was performed. **f,** *In vitro* Rab5 GTPase assay shows that TBC1D5 does not efficiently hydrolyze Rab5. Recombinant Rab5 protein was pre-loaded with GTP. GTP-Rab5 was incubated with various concentration of recombinant TBC1D5. After the reaction, the phosphate release was measured. Three independent experiments were performed. Ordinary one-way ANOVA with Tukey’s HSD was performed. Data are represented as mean ± SEM. ****p < 0.0001, ***p < 0.001, **p < 0.01, and *p < 0.05.

**Extended Data Fig. 3.**
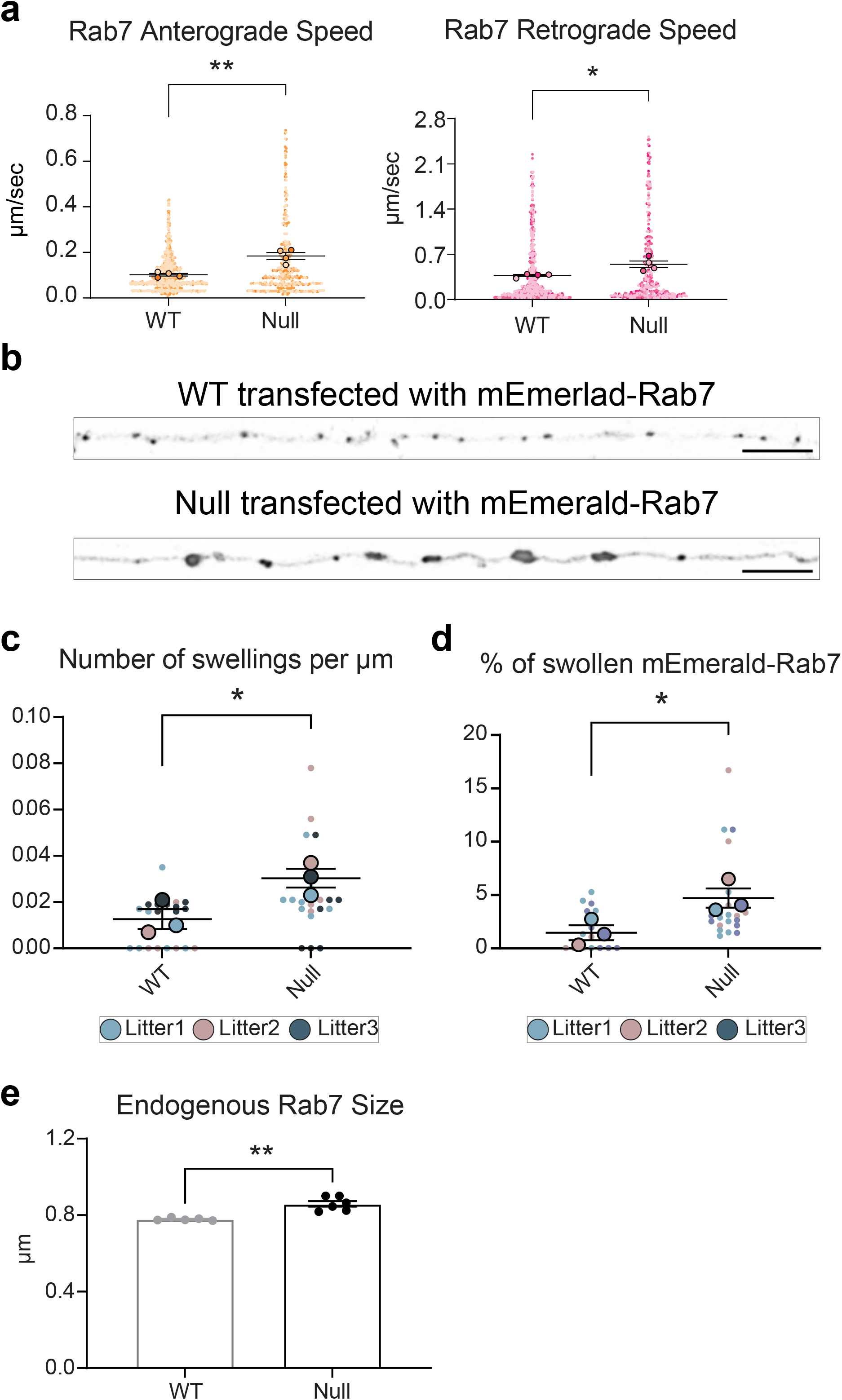
Speed and size of Rab7 endosomes. **a,** Stationary Rab7 endosomes were not included for the calculation. Primary neurons from WT and NHE6-null rats were transfected with mEmerlad-Rab7. 4 independent experiments from 4 different litters were conducted (WT = 5, Null = 5 pups). Two-way ANOVA with Tukey’s HSD was performed. **b,** Representative confocal images of primary neurons from WT and NHE6-null rat brains. Primary neurons were transfected with mEmerald-Rab7 a day prior to the imaging. The size of mEmerald-Rab7 in NHE6-null neurons was larger than that of WT neurons. Scale bar, 5 μm. **c,** Quantification of % swollen mEmerald-Rab7. The number of swollen mEmerald-Rab7 puncta was divided by the total number of mEmerald-Rab7 puncta found within the 50 μm from the AIS. Rab7 endosomes were considered to be swollen if Rab7 puncta’s diameter size is larger than 1.5 μm. **d,** Quantification of the number of swollen mEmerald-Rab7 per μm. The number of swollen mEmerald-Rab7 was divided by 50 μm, to present how many swollen mEmerald-Rab7 exists every μm. **e,** Size of endogenous Rab7 is larger in primary NHE6-null neurons than WT neurons. Data are represented as mean ± SEM. ****p < 0.0001, ***p < 0.001, **p < 0.01, and *p < 0.05.

**Extended Data Fig. 4.**
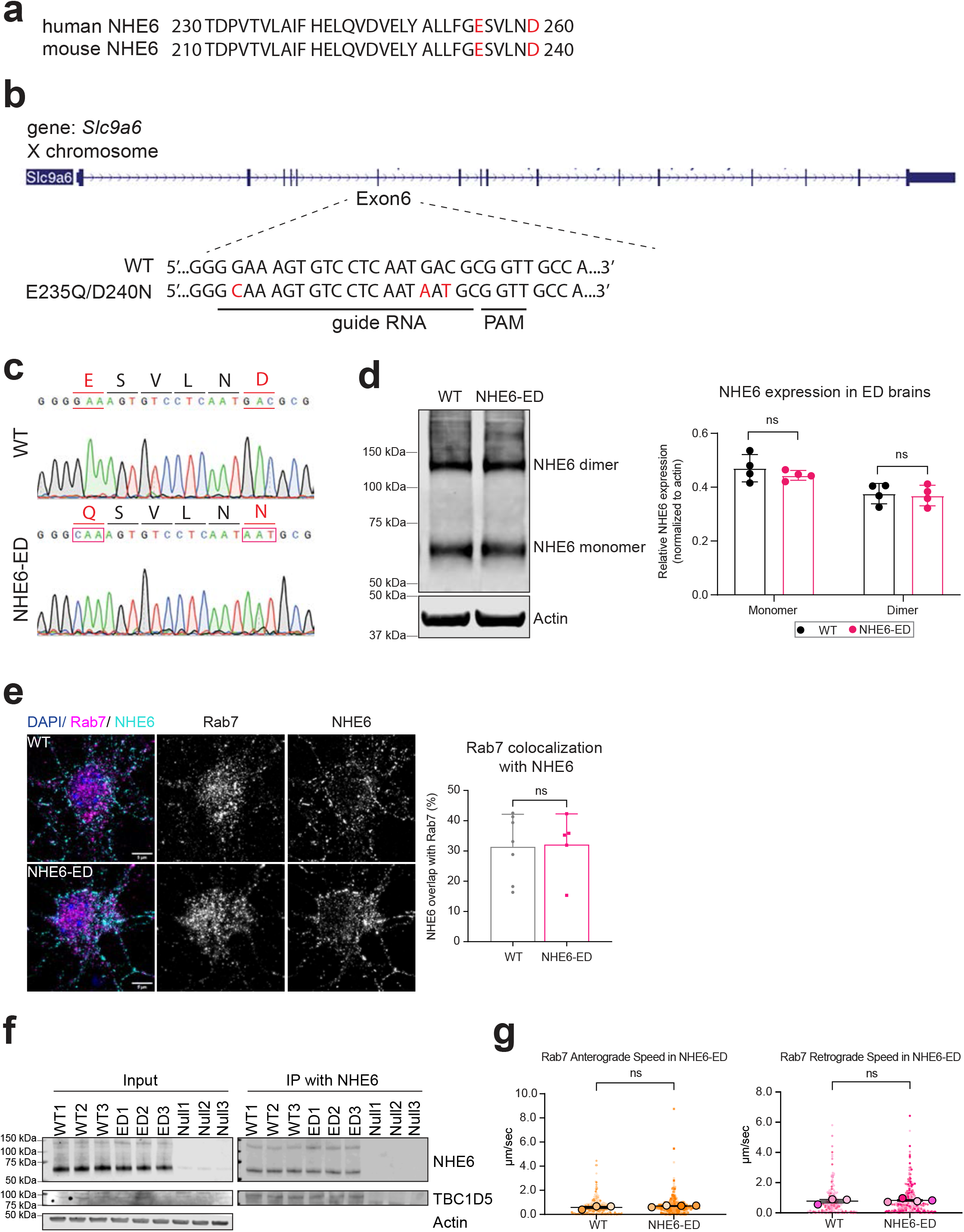
Validation and characterization of the NHE6 exchanger- defective (NHE6-ED) mouse line. **a,** Alignment of amino acid sequences of human mouse NHE6 from a part of the transmembrane domain. The glutamic acid [E] at position 235 in mouse NHE6 corresponds to the glutamic acid [E] at position 255 in human NHE6 (both in red font). The aspartic acid [D] at position 240 in mouse NHE6 corresponds to the aspartic acid [D] in human NHE6 260 (both in red font). **b,** Schematic of the targeted exon of mouse *Slc9a6* (NHE6). The target sequence for CRISPR/Cas9 genome editing is displayed (wild-type (WT), top; NHE6 E235Q/D260N, bottom). Red font indicates the E235Q point mutation (G > C) and the D260N mutations (G>A, C>T). The targe sequences of guide RNA and the protospacer adjacent motif (PAM) are underlined. **c,** Sanger sequences of WT and NHE6-ED mutant mice. Genomic DNA sequence, isolated from mice tail biopsy, presents the sequence in the WT mice and the substitution on the corresponding positions. The presence of the (G > C) point mutation along with (G>A & C>T) mutations were confirmed in samples from NHE6-ED mouse lines. **d,** Western blot of mouse brain lysates from WT and NHE6-ED mutant mice. The expression of NHE6 was detected in brain lysates from both WT and mutant mice for the NHE6-ED mouse line. Quantification of NHE6 densitometry was performed separately for bands at 70 kDa, indicating monomer form of NHE6 and 140 kDa, indicating dimer forms of NHE6. NHE6 expression was normalized to actin. No significant difference in NHE6 levels was detected between WT versus mutant mice. Each dot indicates each animal (WT = 4, Null = 4 pups, n = 10,000-14,000 neurons imaged from each pup). Two-tailed unpaired t-test with Welch’s correction was performed. **e,** Colocalization of Rab7-labeled late endosomes (magenta) and NHE6 (cyan) in primary neurons from littermate WT and NHE6-ED mouse hippocampi. The quantification showed the Mander’s coefficient (% NHE6 overlapping Rab7) from the high-content imaging. No significant difference was detected. Each dot indicates a mean of each pup from 3 different litters (WT = 6, Null = 6 pups, n = 10,000-14,000 neurons imaged from each pup). Two-tailed unpaired t-test with Welch’s correction was performed. Scale bar, 5 μm. **f,** Immunoprecipitation from WT, NHE6-ED and NHE6-null mouse brain lysates shows the interaction of NHE6 with TBC1D5. Brains were immunoprecipitated with anti-NHE6 antibody and immunoblotted against NHE6 and TBC1D5 (n = 3 per each genotype). NHE6-null mouse brains were used as a negative control. **g,** Speed of anterograde and retrograde Rab7 endosomes in primary WT and NHE6-ED neurons. No significant differences were detected between WT and NHE6-ED neurons. Two-tailed unpaired t-test with Welch’s correction was performed. Data are presented as the mean ± SEM. Data are represented as mean ± SEM. ****p < 0.0001, ***p < 0.001, **p < 0.01, and *p < 0.05.

**Extended Data Fig. 5.**
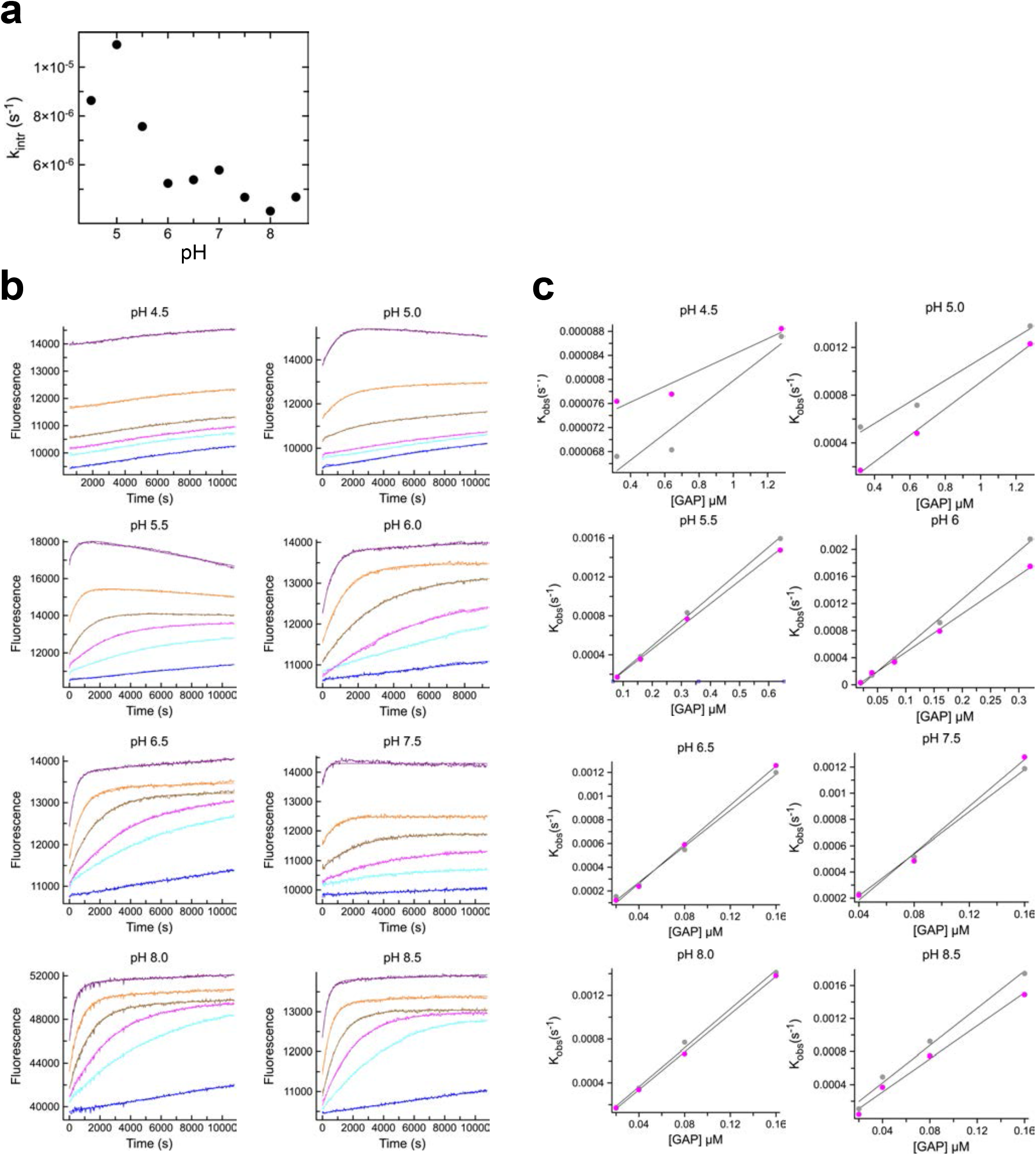
Intrinsic hydrolytic activity and TBC1D5-accelerated hydrolytic activity of Rab7 at pH 4.5-8.5. **a,** Initial rate of intrinsic GTP hydrolysis on Rab7 as a function of pH. In the absence of ceTBC1D5, the rate of intrinsic GTP hydrolysis of Rab7 was very slow (6x10^-6^), and barely changed in the pH range 4.5-8.5. **b,** Intrinsic tryptophan fluorescence time courses of Rab7-GTP hydrolysis in the absence and presence of the *Caenorhabditis elegans* TBC1D5 ortholog (RBG-3 referred as ceTBC1D5) at pH 4.5-8.5. Solid lines are fits of exponential models to the experiment data. This assay was repeated in the pH range 4.5-8.5 to determine kobs. **c,** Fitted kobs for Rab7 as a function of RBG-3 concentration at pH 4.5-8.5. Solid lines are linear fits used to obtain kcat/KM.

**Extended Data Fig. 6.**
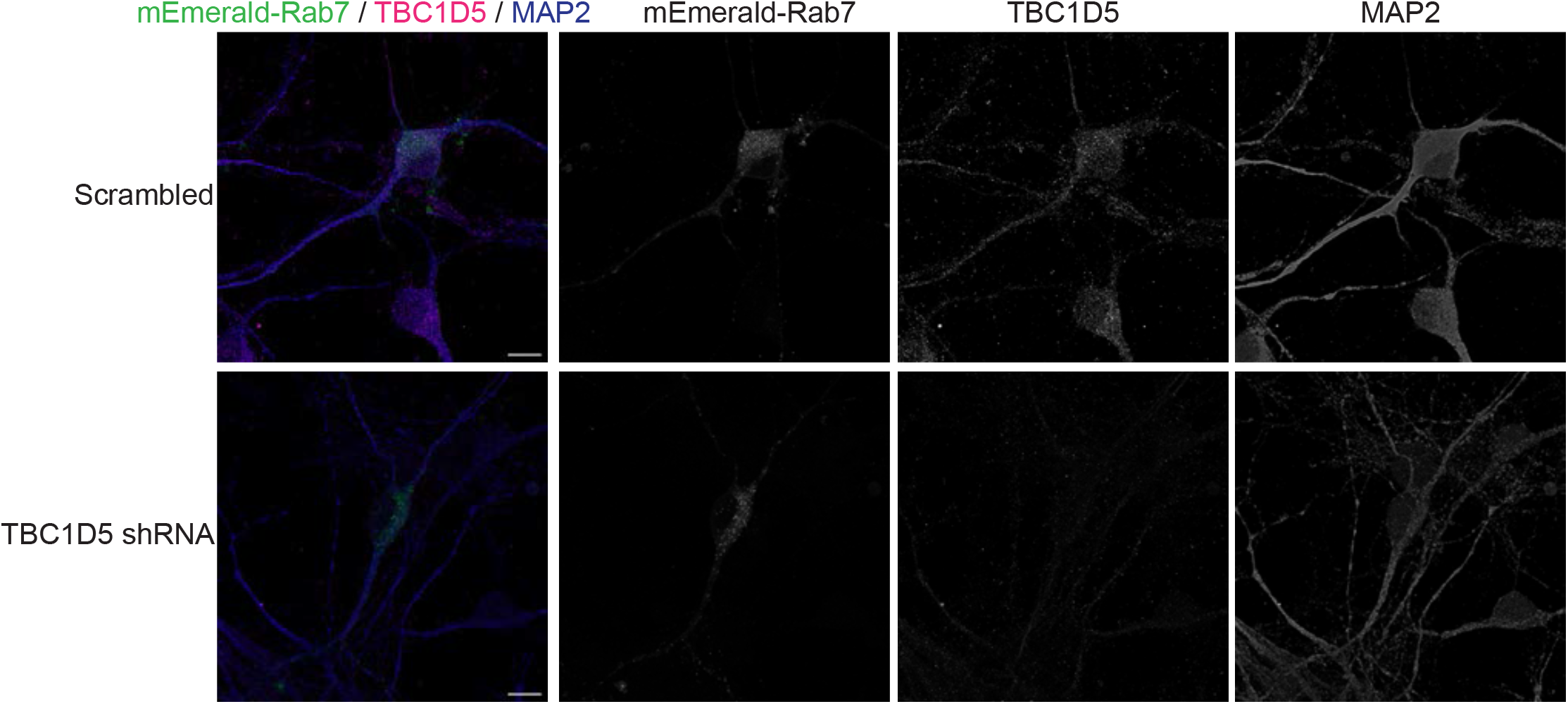
Representative images of TBC1D5 staining after TBC1D5 shRNA transduction. Transduction of TBC1D5 shRNA decreased the expression of TBC1D5, but did not affect the expression of mEmerald-Rab7 in primary neurons, as compared to scrambled shRNA control transductions where there was no effect on TBC1D5. Scrambled shRNA and TBC1D5 shRNA (MOI = 2) were transduced in primary neurons at DIV 2 and incubated for 72 hours. Primary neurons were transfected with mEmerald-Rab7 a day prior to the imaging, and stained with anti-mEmerald (green), TBC1D5 (magenta), and MAP2 (blue; neuronal marker) antibodies. Scale bar, 5 μm.

**Extended Data Fig. 7.**
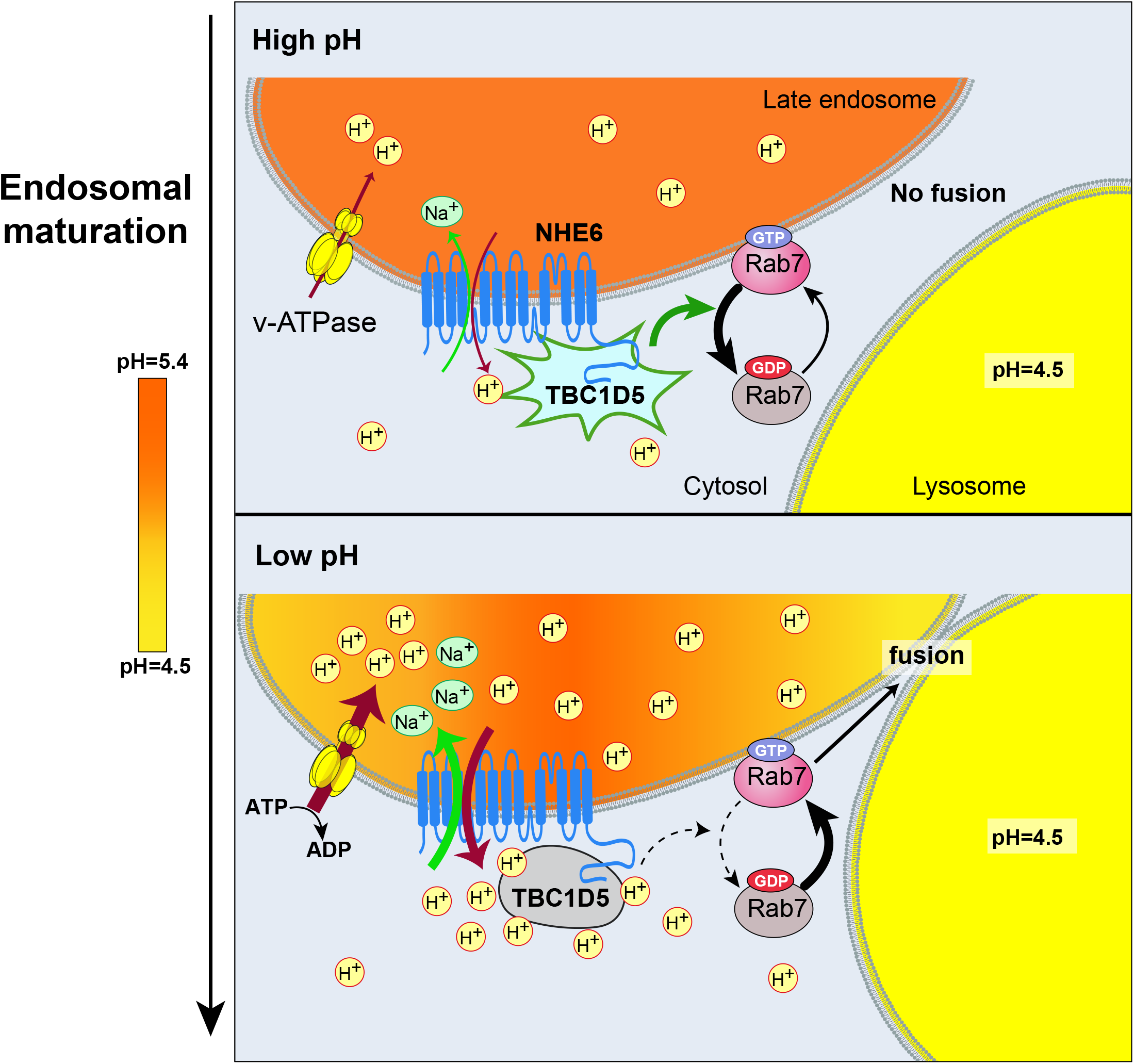
Proposed mechanism by which endosome lumen acidification regulates Rab7 and endosome maturation, a process mediated by proton efflux through CS protein NHE6 and a pH-dependent Rab7 GAP. During endosome maturation, the v-ATPase pumps protons into the endosomal lumen. Early in endosome maturation, as the proton concentration is relatively lower (higher pH) **a,** proton efflux through NHE6 is limited, and TBC1D5 active, accelerating Rab7- GTP hydrolysis. This promotes conversion to inactive Rab7-GDP. Consequently, late endosomes are not efficiently fused with lysosomes. As endosomes mature, the v-ATPase more actively pumps protons into the endosomal lumen. **b,** During this phase, as proton concentration is higher (lower pH), NHE6 leaks protons from late endosomes. As late endosomes gradually acidify, the cytoplasmic microenvironment adjacent NHE6 on late endosomes is progressively acidified. This change in pH inactivates TBC1D5. The GAP activity of TBC1D5 on Rab7 is inhibited, and as a result, Rab7 is converted to the active GTP-bound form, driving a molecular cascade leading late endosomes to fuse with lysosomes.

